# Evolutionary history and genomic vulnerability of the extinct giant deer *Megaloceros giganteus*

**DOI:** 10.64898/2026.05.13.724764

**Authors:** Mikkel-Holger S. Sinding, Marco Gargano, Zihe Li, Emiliano Trucchi, Patrick Arnold, Kevin G. Daly, Valeria Mattiangeli, Sergei Kliver, David Duchene, Daniel Wegmann, Andreas Fueglistaler, Federica Alberti, Greg Gedman, Kathleen Morrill Pirovich, Brandi Cantarel, Iva Kovacic, Joseph Nesme, Lisen Li, Nicola S. Heckeberg, Nigel T. Monaghan, Wilfried Rosendahl, Doris Doeppes, Victoria E. Mullin, Patrícia Pečnerová, Rupinder Kaur, Christian Carøe, Sarah S. T. Mak, Love Dalén, Wen Wang, Eline D Lorenzen, Beth Shapiro, Daniel G Bradley, Michael Hofreiter, M. Thomas P. Gilbert, Michael V. Westbury

## Abstract

The extinct giant deer (*Megaloceros giganteus*) was one of the most striking megafaunal species of the Late Quaternary, distinguished by its enormous palmated antlers reaching up to 3.5 m across, the largest known among both living and extinct cervids. Despite its iconic status, little is known about its genomic history prior to extinction ∼8 thousand years ago (kya). We generated the first nuclear palaeogenomes for *Megaloceros*, represented by nine individuals from Germany (∼40 kya) and Ireland (∼11 kya), mapped to a new chromosome-level reference genome of the fallow deer (*Dama dama*). Phylogenomic analyses placed *Megaloceros* as sister to *Dama* (divergence ∼3.5 Ma) and revealed evidence of gene flow with ancestral *Cervus* lineages. Population analyses identified clear differentiation between German and Irish lineages, with higher genetic diversity in the German individuals. Two genes under strong positive selection, BNIPL and SLC10A7, are associated with apoptosis regulation and skeletal development/bone mineralisation, respectively, and may relate to the species’ large body and antler size. Demographic reconstructions indicate a long-term decline in effective population size, extremely low heterozygosity, little evidence of extensive runs of homozygosity, and an elevated burden of predicted deleterious alleles. Together, these results suggest that *Megaloceros* entered the terminal Pleistocene in a genomically fragile state, offering new insight into the biology and evolutionary legacy of one of the largest and most distinctive cervids that ever lived.

## Introduction

At the end of the Late Pleistocene (∼11.7 thousand years ago; kya) a substantial proportion of the world’s large terrestrial mammals - over 70% of megafaunal genera in some regions - went extinct ^1,2^. Among the most distinctive of these taxa is the giant deer, *Megaloceros giganteus* (hereafter *Megaloceros*), renowned for its enormous palmated antlers reaching up to ∼3.5 m in span, the largest known in any cervid ^3^. Although *Megaloceros* has long held an iconic place in Quaternary palaeontology, key aspects of its evolutionary history, population structure, and genomic biology remain poorly resolved.

Most recent taxonomic syntheses support *Megaloceros* as a single, widely distributed species within Cervinae ^3–7^. However, its precise taxonomic affinities among cervids have been debated for over a century. Based on different combinations of morphological characters and limited molecular data, *Megaloceros* has historically been phylogenetically placed in a variety of positions within Cervinae, including affinities with *Cervus* ^6,8–11^, *Rangifer* ^12^, the damine deer (*Dama*) ^4,6,13–17^, or as the sister taxon to all cervine ^18^. While multiple morphological and mitochondrial studies have increasingly supported an affiliation with the damine group, alternative placements have persisted across datasets and analytical frameworks.

Despite its nickname as the “Irish elk”, *Megaloceros* was not uniquely Irish but occupied a broad Late Pleistocene range across northern Eurasia, with higher abundance in interglacials compared to glacial periods ^19^. Radiocarbon dates on *Megaloceros* remains indicate that it persisted into the Holocene, surviving in western Siberia and eastern Europe until ∼7,660 calibrated years before present (cal BP) ^20^. During climatic fluctuations its range cycled, shrinking during glacial maxima and expanding during interglacials ^19^. Hypotheses on the extinction of *Megaloceros* range from climatic and vegetational shifts affecting forage quality, to intensified hunting pressure by humans and interspecific competition with other cervids ^8,21^.

Nuclear palaeogenomics has transformed our ability to reconstruct the evolutionary histories of extinct species, enabling inference of phylogenetic relationships, past admixture, population structure, effective population size trajectories, and comparative genomic features linked to adaptation or reduced genomic diversity ^22–25^. However, previous studies on *Megaloceros* have been limited to mitochondrial data ^11,20,26^, a single locus which reflects maternal inheritance only, and can be confounded by sex-biased dispersal and gene flow ^25^. Nuclear genomic data are therefore essential for robustly placing *Megaloceros* within the cervid phylogeny, evaluating evidence for historical gene flow, and characterising genome-wide patterns of diversity, inbreeding, demographic history, and candidate adaptive genes. Comparison between late-surviving populations further allows assessment of how genomic diversity and structure differed near the end of the species’ history, and whether long-term demographic trends and signatures of genomic vulnerability were already established.

Here, we generate the first nuclear palaeogenomes for *Megaloceros*, represented by nine individuals: five from the northern Upper Rhine Graben region of Germany dated to ∼40 kya and four from Ireland dated to ∼11 kya, including one high-coverage genome (∼48×) (Fig. 1A). We also produce a new chromosome-level reference genome for the fallow deer (*Dama dama*), facilitating mapping and comparative analyses in the absence of a conspecific *Megaloceros* reference. Using these resources, we investigate the evolutionary relationships of *Megaloceros*, including its phylogenetic placement and historical gene flow among cervids, as well as intraspecific population structure; assess genome-wide patterns of genetic diversity, runs of homozygosity, and the burden of predicted deleterious variation; and infer long-term demographic trends and candidate lineage-specific signals of selection. Together, these data provide a genome-wide perspective on the evolutionary history and genomic characteristics of one of the most remarkable cervids of the Late Quaternary.

**Figure 1:**
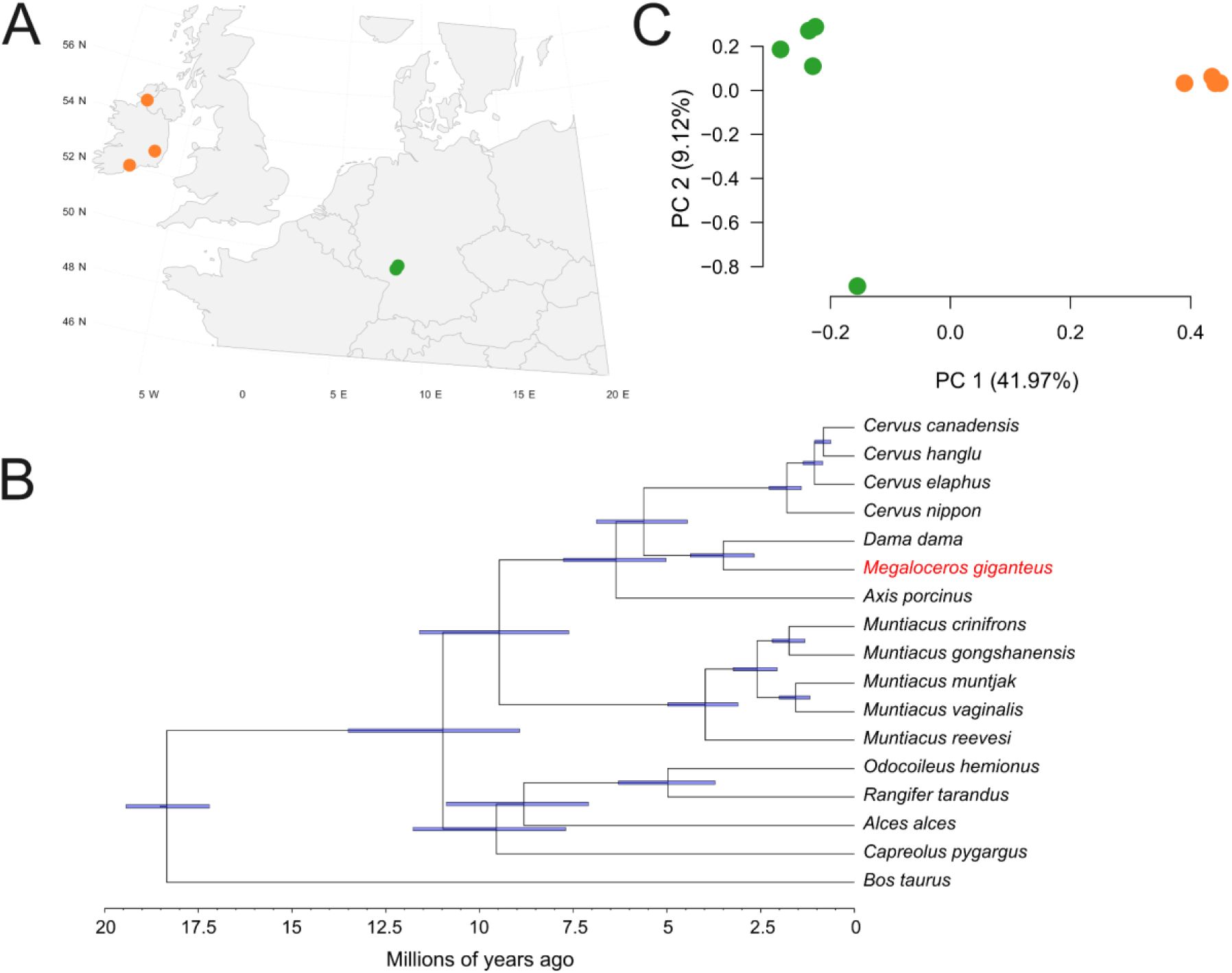
Sample localities and evolutionary relationships. **A)** Map showing the approximate locations of the sites that the *Megaloceros* specimens were obtained from. MG4 is not shown due to lacking locality information. **B)** Fossil-calibrated phylogenetic tree based on the nuclear genomes of *Megaloceros* and selected related species. **C)** Principal component analysis of the nine *Megaloceros* individuals computed using genotype likelihoods. Colours correspond to the sample localities in the map.

## Results

### Megaloceros samples

We analysed nuclear genomic data from nine *Megaloceros* individuals originating from the Late Pleistocene and from two regions in western and central Europe. Four individuals derive from Ireland and date to the Late Glacial (∼12–11 kya cal BP), representing multiple regions of the island (Counties Donegal, Cork, and Laois). Five individuals derive from the northern Upper Rhine Graben region of southwestern Germany (Groß-Rohrheim and Bobenheim-Roxheim) and date to ∼47–37 kya BP (Marine Isotope Stage 3), a period characterised by alternating interstadial and stadial conditions.

### Dama dama assembly

We sequenced 10.37 million (109.7 Gbp, ∼42.6x coverage) PacBio HiFi reads and 1,081.6 million (324.2 Gbp) Arima HiC read pairs from a male European fallow deer (*Dama dama*) from Denmark, and used these to generate a *de novo* diploid chromosome-length genome assembly. We selected fallow deer as previous studies have shown it to be the closest living relative to *Megaloceros*^11^. The obtained (*pseudo*)haplotypes had high QV values (> 65) and had total lengths of 2.38 Gbp (paternal) and 2.72 Gbp (maternal). The paternal (*pseudo*)haplotype was slightly smaller than the genome size estimate of 2.51 Gbp, obtained from the raw HiFi reads (Supplementary Figure S1), whereas the maternal (*pseudo*)haplotype was slightly larger. This difference is probably due to artefacts of assembly in highly repetitive regions like heterochromatin blocks and centromeres. Both (*pseudo*)haplotypes include 34 chromosomal scaffolds (chromosomes) as expected from the karyotype (2n = 68) of *D. dama* ^27^. We observed a one-to-one synteny to the red deer (*Cervus elaphus*), and named the chromosomes accordingly (Supplementary figures S2 and S3). The BUSCO scores reached 97.7% (maternal) and 92.3 % (paternal) for complete genes, indicating a high completeness of the assembly.

### Mapping results

Mapping the *Megaloceros* genomic data to the newly assembled *Dama dama* reference yielded average genome-wide coverages between 0.21–5.64× for the German individuals, 0.4–1.58× for the low-coverage Irish individuals, and one high-coverage Irish genome (MG3, 48×). The mapped reads exhibited characteristic ancient DNA damage patterns, supporting their authenticity (Supplementary Fig. S4). Sequence reads from 16 modern species used for comparative analyses were mapped to multiple reference genomes, producing high-quality datasets with genome-wide coverages of approximately 10–32×. Detailed mapping statistics and reference combinations are provided in Supplementary Tables S1–S2.

### Systematic relationships

Our nuclear phylogenomic analyses recovered the expected topology for the sampled extant cervids (Fig 1B). *Megaloceros* was placed as sister taxon to *Dama dama*, with their divergence time estimated at ∼3.5 Ma (95% highest posterior density, highest posterior density [HPD]: 4.38 – 2.68 Ma). Together, *Megaloceros* and *Dama* formed a sister clade to a clade containing all *Cervus* species included in this study. This clade was sister to *Axis porcinus*. The *Muntiacus* species formed a well-supported monophyletic group, while *Odocoileus hemionus* clustered with *Rangifer tarandus*; together they were sister to *Alces alces*, which was subsequently sister to *Capreolus pygargus*. Our *f*-branch analysis showed several instances of excess allele sharing between lineages, indicating gene flow (Fig 2). When focussing on *Megaloceros*, we see excess allele sharing with each of the *Cervus* species as well as with *Axis porcinus*.

**Fig 2:**
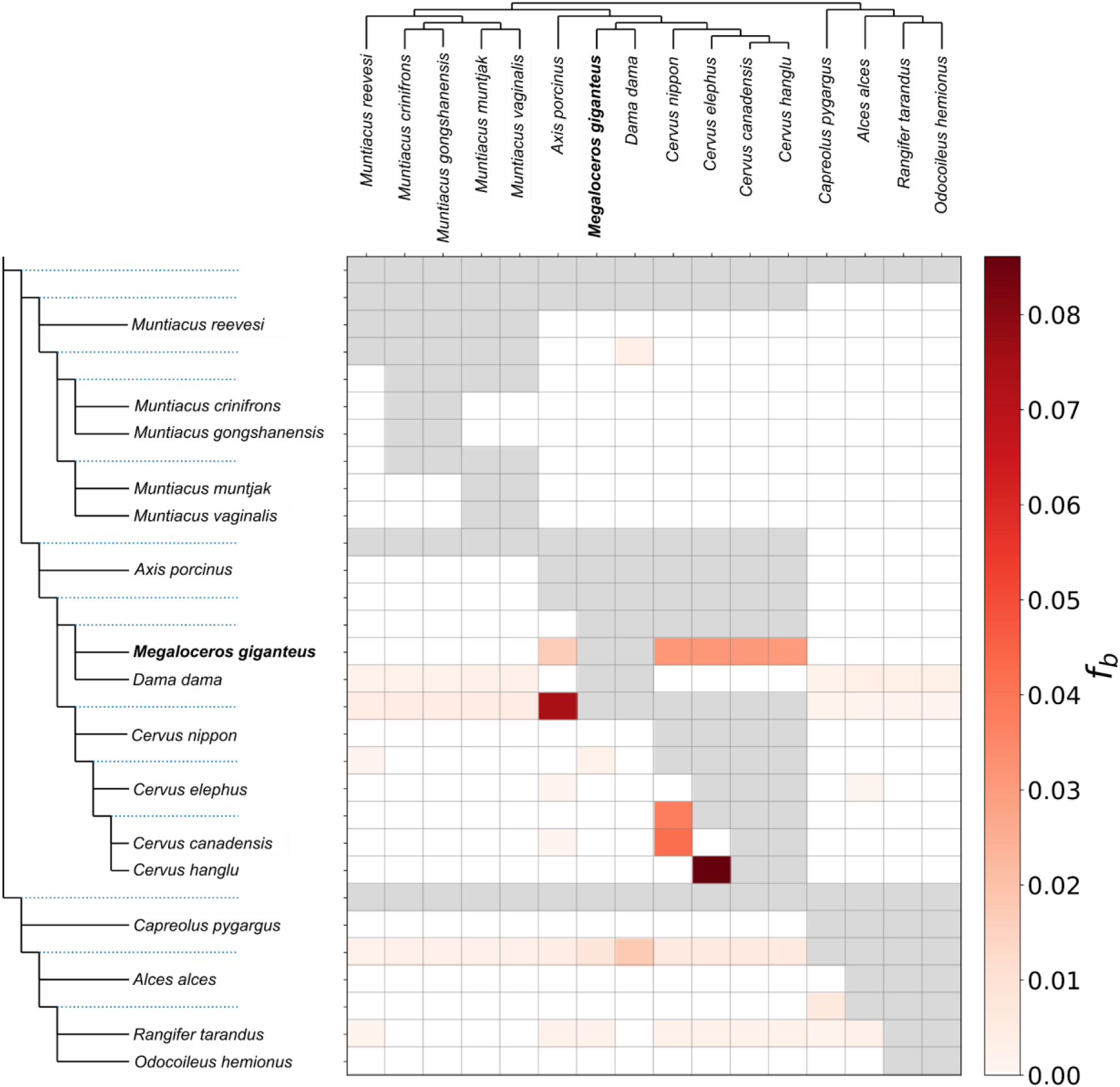
Genome-wide *f*-branch results. The species tree is shown at top and the species tree in expanded form, with internal branches as dotted lines, is shown at left. The values in the matrix refer to excess allele sharing between the expanded tree branch (relative to its sister branch) and the species on the *x*-axis. Grey squares are comparisons that could not be made due to the topological input requirements of the test and its inability to infer gene flow between sister lineages.

### Population relationships

A principal component analysis (PCA; Fig 1C) revealed clear genetic differentiation between the Irish and German *Megaloceros* individuals along the PC1 axis. The PC2 axis explains much less variation within the dataset (9.12% vs 41.97%) and places a single Rhine individual (NK54) as differentiated. A Tracy-Widom test showed that only PC1 could be considered reliable. The differentiation between Irish and German individuals is replicated in our neighbour-joining tree generated from pseudohaploid base calls (Supplementary Fig S5). However, the NK54 individual separated on the PC2 axis does not appear highly differentiated in the tree. Differentiation between the two geographic regions was corroborated by fixation index analyses (Fst = 0.125, p < 2.2 × 10⁻¹⁶).

The mitochondrial genome topology and divergence time estimates closely resemble those from previous analyses of *Megaloceros/Sinomegaceros* mitochondrial data sets, including major lineages (clades 1-5 from Rey-Iglesia et al.^20^ and clades sino2 and sino3 from Xiao et al. ^26^) (Supplementary Fig S6). The deepest divergence splits clade 1 from all other *Megaloceros/Sinomegaceros* specimens and dates to 199.0 kya (95% HPD: 251.2 – 154.4 kya). *Sinomegaceros* forms two lineages (sino2, sino3) that are basal to clade 2-5. In contrast to the nuclear genomic data, Rhine Valley individuals do not form a monophyletic mitochondrial group but are distributed across multiple deeply diverged mitochondrial lineages. NK1 and Meng16 into the early diverging clade 1 and NK54 into clade 3 whereas Meng14 and NK57 form a separate clade that is basal to clade 2-3. Similarly, previously published mitochondrial genomes from the same Irish *Megaloceros* are interspersed within a single Late Glacial clade rather than forming a distinct mitochondrial sublineage.

### Genomic diversity and inbreeding

To estimate a reliable alternative allele frequency for calling heterozygous positions we visualised heterozygous site calls based on the alternative allele frequencies (Supplementary Fig S7). Through this we see a proportionally large number of heterozygous sites with alternative allele frequencies < 0.1 in all individuals analysed, consistent with allelic imbalance likely arising from sequencing error. Transition heterozygous sites (C-T and A-G) retain high values in *Megaloceros* until ∼0.2, likely due to ancient DNA damage. In all cases, heterozygous calls level off at relatively low values until ∼0.25, where they increase again. Therefore, we only considered a site as truly heterozygous if the less frequent of the two alleles occurred at a relative proportion >0.25.

Mean autosome-wide heterozygosity values in the ten cervid species included in this study ranged between 0.004360 and 0.000356 (Fig 3A and Supplementary table S3), estimated using an allele-count–based approach that applies conservative minor allele frequency thresholds to mitigate sequencing errors and ancient DNA damage. Overall, *Dama dama* had the lowest mean level of autosomal heterozygosity (0.000356). However, when using the highest correction value to attempt to correct for inflated heterozygosity values due to reference bias, *Megaloceros* was slightly lower (0.000348). When using a more conservative or no-correction value, *Megaloceros* was second lowest, only to *Dama dama* (0.000568 and 0.000668, respectively).

**Figure 3:**
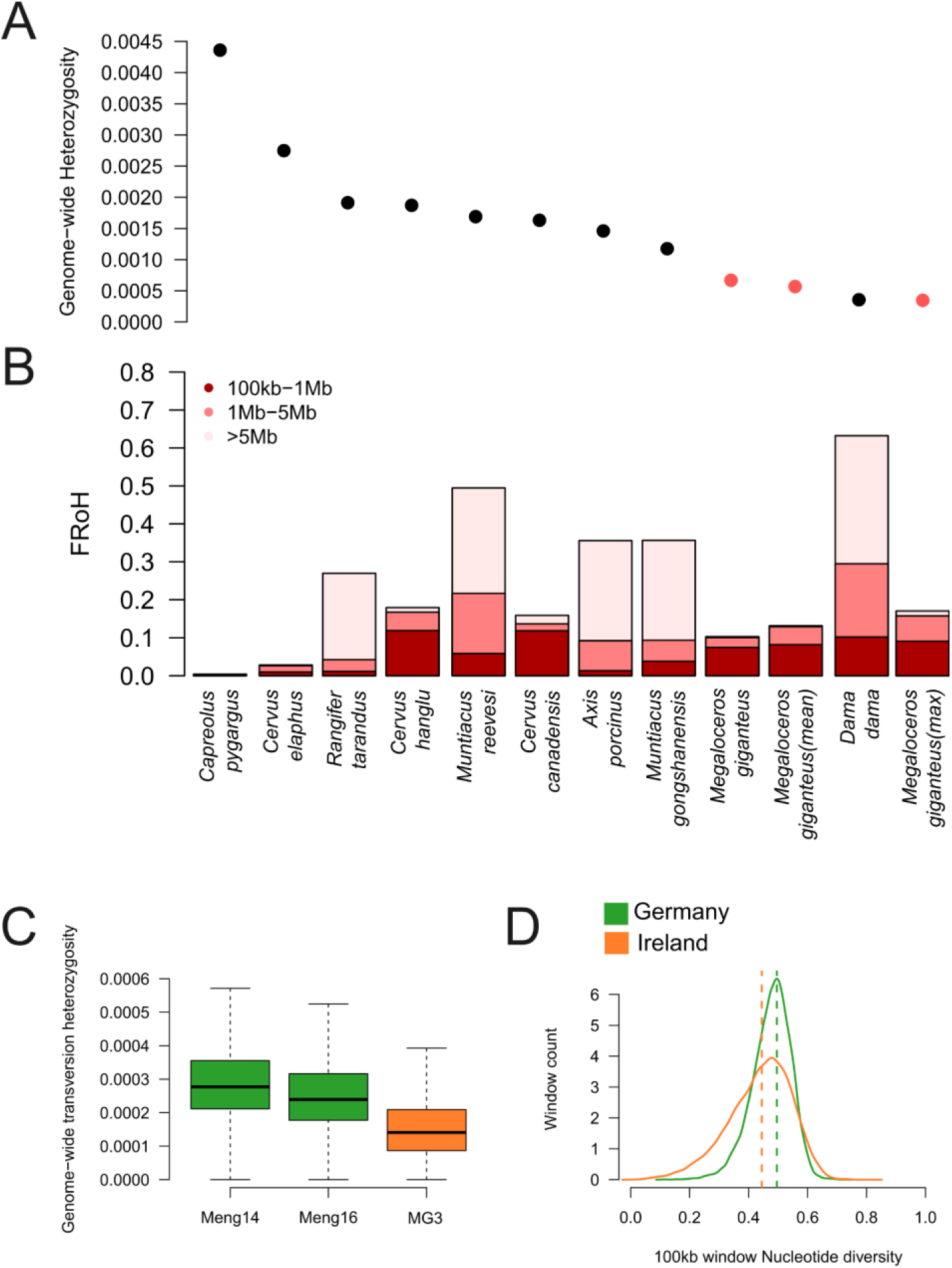
Genetic diversity and inbreeding results within *Megaloceros* and across modern cervids. **A)** Autosome-wide heterozygosity and **B)** one-window bridged runs of homozygosity (a single non-ROH window reclassified as ROH when flanked by ROH windows on both sides) for *Megaloceros* and nine other cervid species. Three values are shown for *Megaloceros*, no correction factor for reference bias, the mean correction bias value, and the maximum correction bias value. Correction values were obtained by mapping extant species to both a conspecific and heterospecific reference genome. **C)** Autosome-wide transversion heterozygosity for the three highest-coverage *Megaloceros* individuals from Ireland (n = 1, red) and Germany (n = 2, blue) in our dataset. MG3 was downsampled to ∼5x to be comparable to the others. **D)** Nucleotide diversity estimates for the two populations (Ireland n = 4; Germany n = 5) counted in 100-kb windows with a total number of 16,346 windows. Dashed lines indicate the mean value.

We observed substantial variation in both the overall proportion and size distribution of ROH across the ten cervid species after applying a strict heterozygosity threshold of <0.0001 to define 100-kb windows as runs of homozygosity (ROH) (Fig. 3B). The highest proportion of long ROH (>5 Mb), indicative of recent inbreeding, was detected in *Dama dama*, followed by *Rangifer tarandus* and *Muntiacus gongshanensis*. In contrast, *Megaloceros* exhibited an intermediate overall fraction of the genome in ROH (FROH) relative to the other species; however, most ROH segments were short (<1 Mb), and no ROH exceeding 5 Mb were detected under this threshold. The lowest FROH values were observed in *Capreolus pygargus*, followed by *Cervus elaphus*.

Mapping modern cervid individuals to a heterospecific reference genome (either fallow or red deer) led to increased autosomal heterozygosity and a decrease of long ROH (especially those >5Mb) in all cases (Supplementary table S3 and Fig S8). While increasing the minimum window heterozygosity threshold for an ROH could recover some of the longer ROH for individuals mapped to the red deer, this was not the case when mapping to the fallow deer (Supplementary table S3). We found no clear correlation between the relative increase in heterozygosity values and the distance to the reference genome (Supplementary Fig S8).

Visual inspection of window-level heterozygosity revealed that, under heterospecific mapping, long ROH were not eliminated but instead fragmented by intermittent windows of elevated heterozygosity (Supplementary Fig. S9). In an attempt to mitigate this effect, we implemented a one-window bridging rule, whereby a single non-ROH window flanked by ROH windows was reclassified as ROH (Supplementary table S4). Bridging partially restored long ROH in modern individuals mapped to heterospecific references. For example, in *Dama dama* mapped to the *Cervus elaphus* reference, raw FROH >5 Mb was reduced to zero, whereas bridging recovered a substantial proportion of long ROH (FROH >5 Mb = 0.103), though still below the conspecific baseline (0.318). A similar pattern was observed in *Cervus canadensis*: when mapped to its conspecific reference, raw FROH >5 Mb was 0.0135 (bridged = 0.0224), whereas mapping to *Cervus elaphus* reduced this to 0.0029 (bridged = 0.0176), again indicating fragmentation under heterospecific mapping and partial recovery with bridging. In contrast, *Megaloceros* (MG3) exhibited no substantial long ROH under either approach (raw FROH >5 Mb = 0; bridged FROH >5 Mb = 0.0026), indicating that the absence of long ROH in *Megaloceros* is unlikely to be solely attributable to reference-induced fragmentation. Despite this, visualising ROH across the genome of *Megaloceros* revealed several instances of ROH (Supplementary Fig. S10)

When only including transversions, we recovered lower levels of heterozygosity and longer ROH but relative comparisons between species remained consistent (Supplementary table S3). We found a clear correlation in heterozygosity between results using all sites and only transversions, including for *Megaloceros*, which has higher rates of transitional (C-T and G-A) errors due to ancient DNA damage (Supplementary Fig S11).

To estimate the relative differences in genetic diversity in each of our respective *Megaloceros* populations (Germany and Ireland), we calculated transversional heterozygosity for the three highest-coverage individuals and nucleotide diversity in sliding windows across the genomes for all nine *Megaloceros* individuals while pooling them into two populations, Ireland and Germany. In both cases we see that the German population exhibited higher mean genetic diversity compared to the Irish population (Fig 3C, 3D and Supplementary table S4).

To complement allele-count–based estimates and to assess the robustness of diversity patterns in lower-coverage individuals, we additionally estimated heterozygosity using a genotype likelihood framework implemented in ATLAS, which explicitly accounts for post-mortem damage and sequencing error through recalibrated quality scores. Heterozygosity was estimated in 5 Mb windows across all autosomes. Estimates were largely insensitive to sequencing depth, remaining stable even when data were down-sampled to 0.1× coverage (Supplementary Fig. S12). Across all analyses, Irish *Megaloceros* consistently showed lower heterozygosity than German individuals. Absolute values varied across individuals, but the relative pattern was consistent across coverage levels and analytical subsets.

### Genetic load

The three Irish samples show more derived homozygous genotypes (realised load) than the four German samples (Fig. 4A), although these estimates are strongly impacted by sequencing coverage (Supplementary Fig S13). To control for the uneven coverage distribution across ancient individuals, we randomly sampled one read at each individual genotype and checked whether it was supporting the ancestral or derived allele. We estimated the allele frequency at each locus and calculated the Rxy, a statistic that compares the relative frequency of derived alleles in one population against another (Supplementary Fig S14), using the Irish samples as the *x* population and the German samples as the *y* population, for each fitness impact class. Using the *modifier* impact class as a neutral background, we calculated the R’_xy_ for *low*, *moderate*, and *high* fitness impact classes. We observe R’_xy_ deviation towards values smaller than 1, indicating lower accumulation of derived alleles at *moderate* and *high* impact variants in the Irish population than expected by drift only (Fig. 4B). This result suggests that selection was efficient at removing deleterious alleles in the Irish population before its extinction.

**Figure 4.**
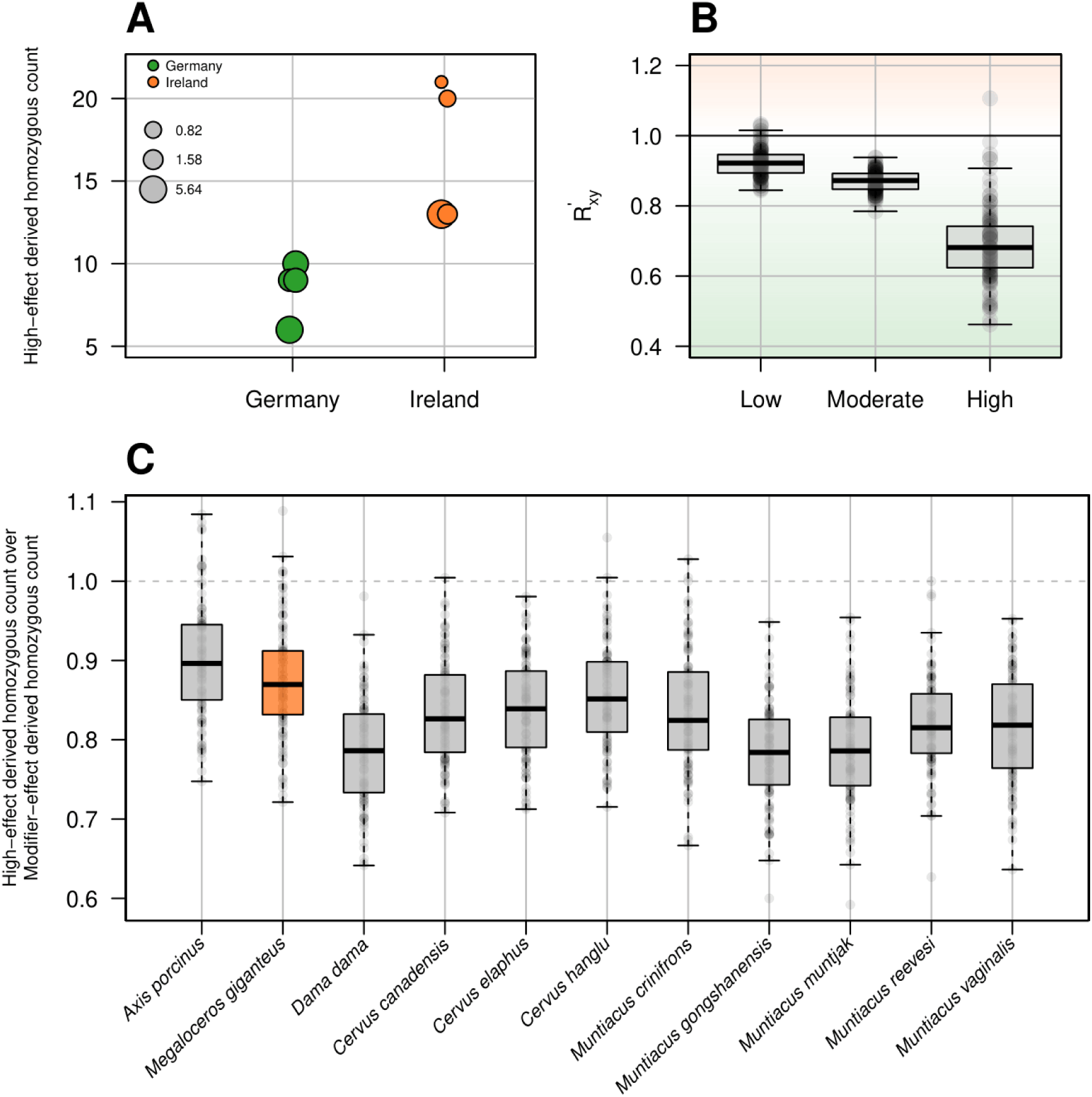
Results of the genetic load analysis. **A)** Number of homozygous genotypes for the derived allele at high-impact sites in German (blue, n = 4) and Irish (red, n = 3) samples. Size of dot is proportional to sequencing coverage. **B)** Distribution of R’_xy_ values calculated between Irish (x) and German (y) populations across 100 pseudo-haploid resampled replicates. The colour gradient highlights the direction and magnitude of the statistic, with increasing red intensity for R’_xy_ > 1 (Irish enrichment) and increasing blue intensity for R’_xy_ < 1 (German enrichment). **C)** Distribution across 11 cervid species of the ratio between the number of homozygous genotypes for the derived allele at high-impact sites and the number of homozygous genotypes for the derived allele at modifier-impact sites, calculated across 100 replicate subsets of 1000 randomly sampled sites per ingroup, using only high-coverage samples.

To evaluate the efficacy of selection acting on the Irish sample in a broader context, we compared the genetic load in the high-coverage *Megaloceros* sample from Ireland (MG3) with the genetic load estimated in high-coverage samples from ten other cervid species. The raw number of high-impact derived genotypes in *Megaloceros* was higher than in all other cervids, except for *A. porcinus* and *M. reevesi* (Supplementary Fig S15). Notably, *Megaloceros* showed one of the highest number of high-impact derived genotypes also when normalizing using the *modifier* impact class as neutral background (Fig 4C). Among all cervids, a similar minor loss of *high*-impact derived homozygous genotypes relative to *modifier*-impact sites was observed in two other species, *A. porcinus* and *C. hanglu* (Fig. 4C).

### Demographic history

Across the last ∼1 Ma, PSMC trajectories indicate species-specific demographic histories rather than a single shared pattern (Fig 5). *Megaloceros* exhibited relatively high Ne at ∼1 Ma but declined steadily from ∼800 kya until ∼150 kya, after which it experienced a brief recovery followed by a sharp contraction toward the end of the Pleistocene. From ∼350 kya onwards, *Megaloceros* maintained lower Ne than any of the other species analysed, with exception of *Dama dama* for a relatively brief period ∼100 kya. Overall, the asynchronous peaks and troughs among taxa reflect distinct demographic responses despite partial overlap in geographic range and ecology, with all species showing late Pleistocene decline, albeit at varying degrees. We see a rapid increase in the last time interval for *Megaloceros* but this may be due to reference bias when mapping to a heterospecific reference genome ^28^.

**Figure 5:**
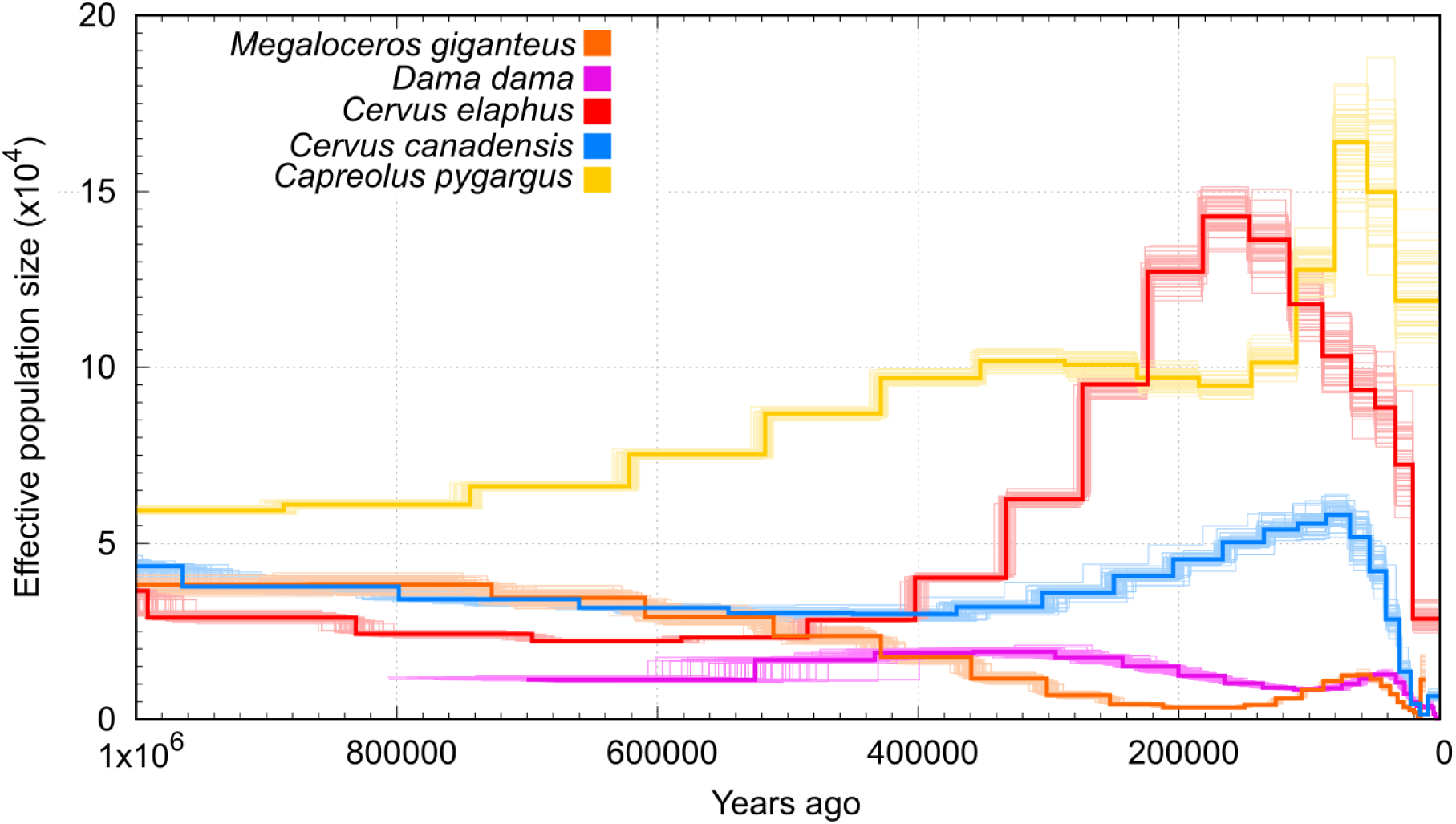
Effective population size of five cervid species calculated using PSMC. Faded lines show the 100 bootstrap replicates produced for each species. Comparative cervid species were selected for their phylogenetic and ecological relevance to *Megaloceros*, including red deer (*Cervus elaphus*; overlapping Eurasian distribution), eastern roe deer (*Capreolus pygargus*; forest–steppe habitats), wapiti (*Cervus canadensis*; large-bodied mixed feeder), and fallow deer (*Dama dama*; closest living relative).

**Figure 6:**
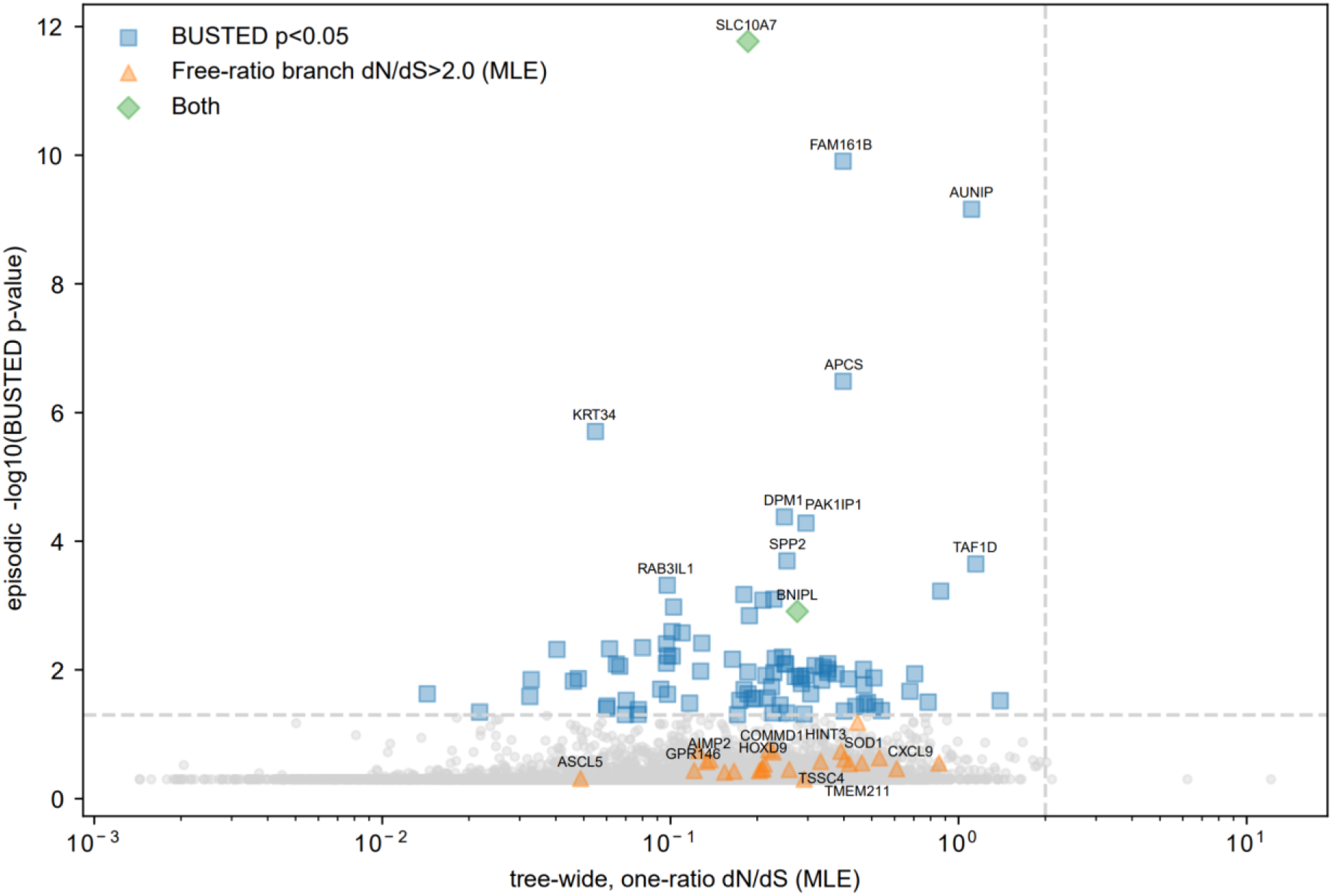
Gene selection analysis of *Megaloceros*. Each point represents one gene, with the x-axis showing the tree-wide one-ratio dN/dS (maximum likelihood estimate, MLE) and the y-axis showing –log10 transformed BUSTED p-values for episodic selection on the *Megaloceros* branch. Grey circles denote genes without significant evidence of selection. Blue squares indicate genes with significant BUSTED support (p < 0.05), orange triangles highlight genes with elevated branch-specific ω values (>2.0), and red diamonds represent genes identified by both tests. Dashed lines indicate thresholds for ω = 2.0 (vertical) and p = 0.05 (horizontal).

### Genes under selection

After whole-genome alignment, we successfully annotated 11,906 ortholog genes relative to the human annotation in the 14 cervid species used in the genes under selection analysis. Using Hyphy framework on these genes, we identified 93 episodic positively selected genes (p<0.05) with the BUSTED model and 26 genes under pervasive positive selection (free-ratio dN/dS > 2) in the *Megaloceros* lineage (Supplementary table S5).

Using Metascape^29^, the 93 genes under episodic positive selection were clustered into a set of significantly enriched functional terms and pathways (q < 0.05), spanning Gene Ontology biological processes as well as KEGG, Reactome, and WikiPathways categories (Supplementary table S6). The strongest enrichments involved growth, tissue remodeling, and metabolic signaling (24 genes across 4 terms), including IGF regulation, post-translational phosphorylation, and Rap1/PI3K-AKT pathways (SPP2, GOLM1, HGF, VAV2, BCAR1). Hemostatic and vascular processes were also enriched (12 genes across 2 terms), including platelet activation and coagulation pathways (F10, KIF1A, DGKD). We additionally observed enrichments in DNA repair and recombination (5 genes, 2 terms; TOP3A, AUNIP, RNF169), fatty acid derivative metabolism (3 genes, 1 term; ACSM2A, THEM5), immune regulation (6 genes across 2 terms; IL23A, IL4I1, ARID5A), and RNA processing/cell junction assembly (6 genes across 2 terms; ACIN1, STRN, AFDN). Together, these enrichments highlight functional divergence in *Megaloceros* in pathways related to growth, metabolism, vascular function, genome maintenance, immune modulation, and cellular signaling.

Among these genes, BNIPL (BCL2/adenovirus E1B 19kDa interacting protein-like) and SLC10A7 (solute carrier family 10 member 7) stood out as robust candidates, simultaneously exhibiting significant BUSTED p-values (p < 0.05), elevated branch-specific dN/dS, and likelihood ratio tests supporting the branch-specific model over the global model.

## Discussion

Our study provides a genome-wide perspective on the evolution and extinction of *Megaloceros*. We confirm its evolutionary placement as the sister taxon to *Dama dama* and detect evidence of gene flow with *Cervus*. We also identify signatures of selection in genes linked to body size and skeletal growth that may have underpinned its iconic gigantism. Furthermore, we uncover genetic patterns consistent with long-term demographic decline, low effective population size, reduced genetic diversity, and elevated genetic load. Together, these findings illuminate both the genetic basis of the evolutionary innovations that defined *Megaloceros* and the demographic and genomic context of its Late Quaternary populations, illustrating how palaeogenomics can refine our understanding of evolutionary history in extinct megafauna.

### Evolutionary history

Our phylogenomic results are in agreement with previous studies ^4,5,17^ using mitochondrial DNA that have placed *Megaloceros* and *Dama* as sister taxa (Fig 1B). Previous mtDNA time-trees have placed the *Megaloceros*–*Dama* split in the late Miocene to early Pliocene, with estimates ranging from ∼5 to ∼10 Ma depending on calibration and dataset ^11,30^. Our nuclear genomic split of ∼3.5 Ma (95% HPD: 4.38 – 2.68 Ma) is considerably younger than this, which may reflect differences in coalescent dynamics between markers, as mitochondrial lineages often trace deeper divergences, whereas genome-wide nuclear data capture a distribution of gene histories around the true species split. Furthermore, by restricting to windows concordant with the species tree, our approach reduces the upward bias from incomplete lineage sorting, yielding a more recent but likely more accurate divergence time ^31^.

Our results also show that the evolutionary history of *Megaloceros* was not a simple bifurcating process (Fig 2). Our finding of excess allele sharing between *Megaloceros* and all four *Cervus* species investigated here could be due to independent gene flow events between *Megaloceros* and each species. However, a more conservative explanation would be ancestral gene flow between *Megaloceros* and the ancestor of the four *Cervus* species included in this study. Given that the divergence between *Megaloceros* and *Dama* is estimated at ∼3.5 Ma, and that in our analysis the *Cervus* crown group diverged ∼ 1.9 Ma (95% HPD: 2.28 – 1.41 Ma), this constrains the timing of the inferred gene flow to between ∼3.5 and ∼1.9 Ma.

After diverging from the *Dama* lineage, *Megaloceros* evolved a highly distinctive phenotype, notably its large body size and massive antlers, traits that have long been hypothesised to be shaped by sexual selection ^3^. Genome-wide tests for positive selection identified multiple genes and pathways potentially associated with these traits. Among the genes inferred to be under selection, BNIPL and SLC10A7 emerged as particularly robust candidates, showing consistent support across multiple metrics, including significant BUSTED tests, elevated branch-specific dN/dS ratios, and improved fit of branch-specific models. BNIPL encodes a BCL2-interacting protein involved in the regulation of apoptosis and cell survival, processes that can influence tissue growth and remodelling during development ^32^. SLC10A7 plays a key role in glycosaminoglycan synthesis and skeletal development, with mutations causing skeletal dysplasia and shortened bones in humans and mice ^33^, making it a plausible contributor to the exceptional skeletal growth characteristic of *Megaloceros*.

Beyond these individual genes, pathway-level enrichment analyses provide broader insight into the molecular processes that may have underpinned the evolution of gigantism and antler development in *Megaloceros*. Genes inferred to be under episodic positive selection (BUSTED) were significantly enriched for pathways involved in growth regulation, tissue remodelling, and metabolic signalling, including IGF regulation and post-translational phosphorylation pathways, which are consistent with selection on rapid skeletal growth and the development of large antlers ^34,35^. These growth-related processes may also relate to the pronounced cranial pachyostosis observed in *Megaloceros* ^16^, a distinctive feature involving substantial thickening of the skull bones. Enrichment of hemostatic and platelet activation pathways may reflect adaptations required to maintain vascular integrity during rapid seasonal tissue expansion, such as antler growth. Additional enrichments in DNA repair and recombination pathways suggest selection on genome maintenance processes, potentially supporting longevity or resilience to physiological stress ^36^. Together, these results suggest that positive selection in *Megaloceros* involved both individual genes with strong signatures of selection and broader functional categories related to growth, metabolism, vascular function, and cellular maintenance.

### Population structure

Previous mitogenomic studies of *Megaloceros* documented high mitochondrial lineage diversity and weak phylogeographic structure across Late Pleistocene Eurasia, with deeply diverged mitochondrial lineages often occurring in close geographic and temporal proximity ^20,26^. Our nuclear genomic results refine this picture by revealing clear nuclear genomic differentiation between Irish and German *Megaloceros*, despite the persistence of multiple mitochondrial lineages within each region. In Germany, individuals that cluster together in nuclear analyses are distributed across several deeply diverged mitochondrial clades, while previously published Irish mitochondrial genomes fall within a single Late Glacial clade, but do not form a distinct monophyletic group (Supplementary Fig S6). This mitochondrial–nuclear discordance indicates that mitochondrial lineage structure was largely decoupled from late-stage nuclear population structure, consistent with the retention of deep mitochondrial coalescent diversity and/or gene flow across populations.

The genomic patterns observed in Ireland should also be interpreted in light of its Late Glacial geographic context: although no substantial, enduring land bridge existed after ∼15 ka, connectivity to mainland Europe likely declined progressively, with full insularity established by ∼10–9 ka ^37^. Although *Megaloceros* ranged widely within the island during the Late Glacial^38^, population size and external connectivity were likely limited following recolonisation, a demographic context expected to enhance the effects of drift and founder events. In contrast, the northern Upper Rhine Graben region occupied a central position within the continental European landscape and likely remained better connected to surrounding *Megaloceros* populations, facilitating ongoing gene flow and the maintenance of higher nuclear genetic diversity. Together, these results suggest that differences between Irish and German *Megaloceros* populations reflect a combination of insularity and spatial connectivity, while reinforcing earlier conclusions that mitochondrial phylogeography alone provides an incomplete representation of population structure in this species.

### Genomic health

Across the cervid panel analysed here, *Megaloceros* exhibited among the lowest levels of autosomal heterozygosity, second only to the Danish *Dama dama* individual, depending on the degree of reference-bias correction applied (Fig. 2A). While low genetic diversity is often associated with small population size ^39^, present-day *D. dama* populations, including those in Denmark where the analysed genome is from, reflect extensive human-mediated translocations ^40,41^ which may have led to increased inbreeding and reduced diversity. Demographic reconstructions of *Megaloceros* indicate that this reduced diversity reflects a long-term history of relatively low effective population size followed by a pronounced decline beginning ∼60 kya (Fig. 5). In line with this trajectory, *Megaloceros* also carries a high burden of predicted deleterious alleles relative to modifier sites (Fig. 4), consistent with reduced efficacy of purifying selection under prolonged demographic contraction.

Temporal comparisons between the older German (∼40 kya) and younger Irish (∼12 kya) individuals provide additional nuance in patterns of deleterious variation. Raw counts of homozygous derived genotypes at high-impact sites are higher in the Irish individuals, but these estimates are highly sensitive to sequencing coverage and should be interpreted with caution. Using the non-normalized Rxy statistic, we detect no clear difference between the Irish and German populations at high-impact sites (Rxy ≈ 1), suggesting that any apparent increase is marginal or absent. In contrast, when using R′xy, which normalizes high-impact sites by expectations from modifier sites, high-impact variants show values below 1, indicating fewer derived alleles than expected under drift alone in the Irish population. This pattern is consistent with continued purifying selection acting against strongly deleterious mutations during demographic contraction, resulting in partial purging rather than a simple accumulation of genetic load. Similar dynamics have been documented in other Late Quaternary megafauna, including the woolly mammoth, where temporally structured genomes revealed progressive purging of highly deleterious variants prior to extinction ^23^. While our dataset is limited to two time points and geographically distinct populations, the direction of the signal is comparable.

Placing these genetic load patterns in a broader interspecific context further clarifies their interpretation. When compared to modern cervids with varying demographic histories, the Irish *Megaloceros* genome exhibits one of the highest burdens of homozygous derived alleles at high-impact sites, closely matching values observed in *Axis porcinus* (Hog deer) and *Cervus hanglu yarkandensis* (Yarkand deer) (Fig 4), two taxa currently facing conservation challenges ^42^. In both cases, the ratio of high-impact to modifier-derived homozygous genotypes approaches neutrality, suggesting limited purging relative to drift. Thus, although some signal of purging may be detectable within *Megaloceros*, its overall genetic load remained elevated relative to most extant relatives, implying a reduced efficiency of purifying selection.

Patterns of genomic diversity also differ between the two sampled regions. The German individuals display higher nuclear diversity than the Irish individuals (Fig. 2C,D), consistent with their temporal position near the onset of the inferred N_e_ decline and with likely greater spatial connectivity within continental Europe. In contrast, the Irish population occupied a geographically constrained setting during the Late Glacial. Radiocarbon-dated fossil occurrences from Ireland show a pronounced absence of Megaloceros during the Last Glacial Maximum, followed by reappearance in the Late Glacial interstadial, suggesting local extirpation and subsequent recolonisation of the island ^43,44^. Such recolonisation likely occurred within a limited temporal window and may have involved relatively few founders. Although *Megaloceros* ranged widely across Ireland during the Late Glacial ^38^, connectivity to mainland Europe was likely reduced during this period and declined further into the early Holocene, constraining effective population size and amplifying the effects of drift.

Inference of recent inbreeding from genomic data of extinct species is complicated by the absence of a conspecific reference genome. When mapped to the *Dama dama* assembly, *Megaloceros* genomes do not exhibit extensive long (>5 Mb) runs of homozygosity (ROH). Analyses of modern cervids mapped to heterospecific references demonstrate that reference divergence fragments long ROH and inflates apparent heterozygosity ^28^ (Supplementary Table S3). Specifically, mapping modern genomes to a divergent reference reduces the apparent fraction of long ROH, while implementation of a one-window bridging rule partially restores these segments, although not to the conspecific baseline. These results confirm that heterospecific mapping can artificially interrupt long homozygous tracts. However, even after applying this correction, the high-coverage Irish *Megaloceros* individual shows minimal long (>5 Mb) ROH, indicating that the absence of extensive long tracts is unlikely to be solely attributable to reference-induced fragmentation. Although ROH-based inference remains conservative under heterospecific mapping, the overall pattern is more consistent with limited recent close-kin mating than with pervasive severe inbreeding. A comparable genomic signature was reported for the extinct blue antelope (*Hippotragus leucophaeus*), which similarly persisted for hundreds of thousands of years at low effective population size and low diversity, yet showed little evidence of recent inbreeding ^45^. Together, these findings suggest that long-term small population size does not inevitably generate extensive long ROH, and that behavioural structure may partially mitigate the genomic consequences of demographic decline.

Independent osteological evidence supports elevated developmental instability in Irish *Megaloceros*. A high incidence of cervical ribs and other skeletal abnormalities has been documented in Late Pleistocene Irish *Megaloceros* ^46^, phenotypes commonly associated with inbreeding in mammals. Comparable data from continental populations are lacking, preventing assessment of whether such traits were species-wide or regionally amplified. Taken together, the genomic and osteological evidence indicates that the Irish population experienced substantial demographic constraint and elevated genetic load in its final millennia, while continental populations likely retained higher diversity through continued spatial connectivity.

### Regional persistence and disappearance

Based on fossil chronology, the disappearance of *Megaloceros* was spatially and temporally heterogeneous. While the species vanished from much of western Europe by the end of the Late Glacial, it persisted into the early Holocene across parts of eastern Europe and western Siberia, with overlapping late survival dates recorded in both regions^21^. This staggered pattern indicates that regional environmental and ecological conditions strongly influenced population persistence, rather than a single, synchronous extinction event.

Our genomic results document long-term demographic decline, low genetic diversity, and an elevated burden of predicted deleterious variation in western and central European populations of *Megaloceros*. However, these features cannot be directly linked to the timing or causes of regional population loss, and genomic data are not available from the late-surviving Siberian populations. Although Siberia also experienced pronounced climatic and environmental change during the terminal phase of the Late Glacial, the ecological consequences of these changes differed regionally, likely allowing suitable habitats for *Megaloceros* to persist in some areas after populations had disappeared elsewhere. As such, regional persistence is best interpreted in ecological rather than genomic terms.

In western Europe, including Ireland, *Megaloceros* populations occupied landscapes undergoing rapid environmental change during the terminal phase of the Late Glacial. In Ireland, securely dated material indicates absence across the Last Glacial Maximum followed by recolonisation during the Late Glacial interstadial, likely within a relatively narrow temporal window and involving limited population sizes^21^. At the time of occupation of our samples (∼12–11 ka), Ireland was becoming increasingly isolated from mainland Europe due to rising sea levels, with full insularity established shortly thereafter ^37^. This combination of recent colonisation, reduced connectivity, and ongoing environmental change would have constrained demographic expansion.

Viewed more broadly, the staggered disappearance of *Megaloceros* mirrors patterns documented in other Late Quaternary megafaunal taxa, where regional populations persisted in ecological refugia long after extinction elsewhere^47^. Comparable asynchronous survivals have been described in South America and other regions, and have been framed within the “Broken Zig-Zag” model^48^, in which megafaunal populations repeatedly persisted or rebounded in suitable regions before eventual loss. In this context, the decline of *Megaloceros* is best understood as a geographically structured process shaped by regional demographic and environmental histories, with genomic data providing insight into long-term population characteristics rather than a singular explanatory mechanism.

### Limitations

This study has several limitations that should be considered when interpreting the results. No conspecific reference genome is available for *Megaloceros*, and although we mitigated this by generating a high-quality *Dama dama* reference genome and explicitly evaluating the effects of heterospecific mapping, reference divergence likely leads to underestimation of runs of homozygosity and complicates inference of recent inbreeding. Our palaeogenomic sampling is also temporally and geographically limited, with nuclear genomic data available from only two regions (∼40 kya Germany and ∼12 kya Ireland) and no genomic data from late-surviving Siberian populations, restricting inference of species-wide spatiotemporal demographic and genomic variation. In addition, several ancient genomes are of low-to-moderate coverage (<5x), which constrains resolution for analyses sensitive to genotype uncertainty, including fine-scale inbreeding and selection. Finally, while genomic patterns can be placed in a broader environmental and demographic context, direct tests of causal relationships between genomic variation, ecological change, and population persistence are beyond the scope of this study.

## Methods

### *Megaloceros* regional information

#### Ireland

The Irish *Megaloceros* individuals analysed here derive from Lateglacial deposits dated to ∼12–11 kya cal BP, spanning the terminal Allerød interstadial and the onset of the Younger Dryas. Although older direct radiocarbon dates have been reported for Irish *Megaloceros*, the securely dated record is dominated by this Lateglacial interval ^43,44^. During this period, Ireland supported open grassland and sparsely wooded habitats suitable for large grazers. Stable isotope data and spatial distribution of fossil finds indicate that *Megaloceros* ranged widely across the island during the Lateglacial, consistent with an island-wide population ^38^.

#### Upper Rhine Graben, Germany

The *Megaloceros* specimens from southwestern Germany (Groß-Rohrheim and Bobenheim-Roxheim) derive from fluvial gravel deposits of the Mannheim Formation within the northern Upper Rhine Graben. This rift basin acted as a major sediment trap throughout the Quaternary ^49^, preserving extensive Late Pleistocene faunal assemblages ^50^. Recent work on associated taxa, including *Hippopotamus amphibius* and *Mammuthus primigenius*, `demonstrates that the faunal remains can be assigned to Marine Isotope Stage 3 (`∼`60–30 ka) and record alternating cold- and warm-adapted faunal communities` ^51^. These findings indicate that the Upper Rhine Graben served as a localized temperate refugium during interstadial phases of MIS 3, allowing temporary persistence of species typically associated with milder climates, including *Megaloceros*. We confirmed this by directly radiocarbon dating all of the *Megaloceros* fossils used for ancient DNA analysis at the Curt-Engelhorn-Center Archaeometry (CEZA) at the Reiss-Engelhorn-Museen in Mannheim ^52^; Table S1; for details see ^51,53^)

Uncalibrated radiocarbon dates from all specimens included in this study were calibrated using Calib v8.10 ^54^ using the IntCal20 calibration curve.

### *Megaloceros* genetic data generation

#### Irish specimens

Petrous bones were sampled in a dedicated ancient DNA facility at Trinity College Dublin, Ireland, following standard protocols ^55^. Blank tubes were included as controls at each step. For each sample, approximately 120 mg of bone powder was subject to DNA extraction initially described by ^56^ and later modified ^57,58^. 1 ml lysis buffer (0.02 M Tris-HCl; 1.7% SDS; 0.45 M EDTA; 0.65 U Proteinase K) was transferred to each supernatant tube, the tube vortexed, and incubated under constant rotation (700 rpm) for 24 hours at 55 ℃, and then for 24 hours at 37 ℃. At the end of the incubation step, tubes were centrifuged at 13,000 rpm for 10 minutes. We removed the supernatant without disturbing the remaining undigested bone powder. We transferred approximately 1 mL of supernatant to an Amicon® Ultra Centrifugal Filter column and added 3 mL of EDTA to purify and reduce the volume of the extract. We centrifuged the columns at 2,500 rpm for 15 minutes, added a second 3 mL aliquot of EDTA, and centrifuged again for 20–25 minutes until approximately 100 µL of extract remained. We then purified the concentrate using Qiagen MinElute columns according to the manufacturer’s instructions and eluted the DNA in 40 µL of EBT buffer. Purified DNA was then subject to dsDNA library construction ^59^ using 16.25 μL starting DNA; a subset of starting DNAs were first treated with Uracil DNA-glycosylase (5 μL at 37℃ for 3 hours). The resulting libraries were amplified using AccuPrime Pfx Polymerase (Invitrogen), and purified using Qiagen MinElute columns following manufacturer’s instructions. Amplified libraries were sequenced on a HiSeq platform (Macrogen, Seoul) and a NovaSeq 6000 platform (Trinseq, Dublin).

#### German specimens

All pre-amplification steps were carried out in the dedicated ancient DNA facilities at the University of Potsdam, including negative controls for both extraction and library preparation. ∼50 mg of bone powder per sample were collected using a Dremel Fortiflex (9100-21) and a 2.4- to 2.8-mm-diameter drill bit. DNA was extracted following a protocol optimized for highly fragmented DNA ^60^. A total amount of 13 ng of DNA for each extract was initially treated with USER enzyme for 15 minutes at 37 °C (modified from Meyer et al ^61^) to remove uracil residues resulting from cytosine deamination. The USER-treated extracts were then converted into single stranded libraries using the protocol described in Gansauge et al ^62^. The resulting libraries were then amplified and dual-indexed. PCR cycles for amplification were determined in advance using qPCR analysis of the unamplified library. Concentration and length distribution were determined using Qubit 2.0 and 2200 TapeStation (Agilent Technologies), respectively. The single-stranded libraries were either sequenced on an Illumina NextSeq500 system ^63^ at the University of Potsdam in 75bp single-end mode or on an Illumina NovaSeq6000 system at the SciLifeLab, Stockholm, in 100bp paired-end mode.

#### Modern species’ genetic data generation

### *Dama dama* de novo chromosome level genome assembly

#### Sample

Liver, kidney and muscle samples were collected on the 5th of March 2023, from a young but adult male, from a fenced population in Rødvig, Denmark. Fallow deer (*Dama dama*) are a non-native species in northern Europe, including Denmark, where populations originate from historical human-mediated introductions ^64^. The animal was taken down as part of harvest of surplus animals, where the meat is followingly sold for consumption, we took advantage of the scheduled culling to secure the samples.

We extracted high–molecular-weight (HMW) DNA from 25 mg of kidney and muscle tissue from the *Dama dama* reference individual using the MagAttract HMW DNA Kit (Qiagen) following the manufacturer’s protocol. We quantified DNA concentrations with the Qubit dsDNA High Sensitivity (HS) Assay Kit (Thermo Fisher Scientific) and assessed fragment size distributions on a Femto Pulse System (Agilent). We diluted the HMW DNA to ∼30 ng/μL and sheared it to an average fragment size of 12–20 kb (speed setting 31) using a Megaruptor 3 (Diagenode).

We prepared two PacBio HiFi libraries (one from kidney and one from muscle DNA) with the SMRTbell Prep Kit 3.0 (Pacific Biosciences). We verified library concentration and fragment size using the Qubit dsDNA HS Assay and the Femto Pulse System with the 165 kbp Genomic DNA assay (Agilent). We bound sequencing primers and polymerase using the PacBio Binding Kit 3.0 and sequenced four SMRT Cell 8M wells on a PacBio Sequel IIe instrument. We retained only HiFi reads with at least four passes for downstream analysis.

We also prepared Illumina short-read libraries from the same individual using the Revelo DNA-seq Enzymatic Kit for MagicPrep NGS (Tecan) and sequenced them on an Illumina NovaSeq 6000 platform with 150 bp paired-end reads.

To generate Hi-C data, we crosslinked liver tissue (∼270 mg per sample, two samples total) following the protocol of Foissac *et al.* (2019) with minor modifications. We pelleted the cells, stored them at −80 °C, and used approximately 500 ng of crosslinked DNA per library with the Arima High Coverage Hi-C Kit (Arima Genomics, Cat. no. A101030). We quantified DNA on a Qubit 3 fluorometer (dsDNA HS Assay) and sheared proximally ligated molecules to ∼500 bp using a Covaris LE220-plus ultrasonicator. We prepared two Hi-C libraries with the Arima Library Prep Module (Arima Genomics), verified concentration (Qubit dsDNA HS) and fragment size (Agilent 2100 Bioanalyzer, High Sensitivity DNA Kit), and sent the libraries to Novogene for sequencing on an Illumina NovaSeq 6000 platform (150 bp paired-end reads). We generated a chromosome-length diploid genome assembly using the AssemblyBrute pipeline commit 774cb422f4f7f24751b103a62f0b200d341c1151^65^. A detailed description of involved tools ^66–89^ was provided in Kliver et al, 2025^90^. Briefly, the assembly procedure included generation of phased contigs from PacBio HiFi and HiC reads, purging haplotypic duplications, HiC-scaffolding, manual curation, and gap closing. Chromosomal scaffolds were arranged to have the same orientation in both (*pseudo*)haplotypes and were named according to synteny to the *Cervus elaphus* assembly (GCA_910594005.1) ^91^.

#### Modern comparative data

We downloaded raw Illumina sequencing data for a single representative individual from each of 14 extant deer species: *Alces alces* (Eurasian elk), *Axis porcinus* (hog deer), *Capreolus pygargus* (eastern roe deer), *Cervus canadensis* (wapiti), *Cervus elaphus* (red deer), *Cervus hanglu yarkandensis* (Yarkand deer), *Cervus nippon* (sika deer), *Muntiacus crinifrons* (black muntjac), *Muntiacus gongshanensis* (Gongshan muntjac), *Muntiacus muntjak*, *Muntiacus reevesi*, *Muntiacus vaginalis*, *Odocoileus hemionus* (mule deer), and *Rangifer tarandus platyrhynchus* (Svalbard reindeer), and the domestic cattle (*Bos taurus*) as an outgroup (Supplementary Table S2). Publicly available reference genome assemblies were downloaded for a subset of eight of these species (*Axis porcinus*, *Capreolus pygargus*, *Cervus canadensis*, *Cervus elaphus*, *Cervus hanglu yarkandensis*, *Muntiacus gongshanensis*, *Muntiacus reevesi*, and *Rangifer tarandus platyrhynchus)*.

#### Autosome identification

We found putative sex chromosome scaffolds in scaffold-level genome assemblies by aligning them to the cow X (Genbank accession: CM008168.2) and human Y (Genbank accession: NC_000024.10) chromosomes. We performed the alignments using satsuma synteny v2.1 ^92^ with default parameters.

### Data processing

#### Nuclear genomes

For all individuals, we removed Illumina adapter sequences, low-quality reads (mean q <25), short reads (<30bp), and merged overlapping read pairs with Fastp v0.23.2 ^93^. For data sequenced using SE chemistry, we skipped the merging step. We mapped the processed reads (merged or SE only for the ancient individuals) to the specified reference genome using Burrows-wheeler-aligner (BWA) v0.7.15 (41) and either utilising the aln algorithm, with the seed disabled (-l 999), maximum number of gap opens set to 2 (-o 2) and a relaxed mismatch parameter (-n 0.01) for the ancient individuals or the mem algorithm with default parameters for the modern individuals. We parsed the alignment files and removed duplicates and reads of mapping quality score <30 using SAMtools v1.6 (42). We checked for ancient DNA damage patterns of the ancient individuals using the estimateErrors task within ATLAS commit 7b82b51b ^94^.

To accommodate the differing requirements of downstream analyses, we employed multiple reference genome mapping strategies. Analyses requiring direct comparison of homologous sites across species (phylogenomic inference, gene flow analyses, and genetic load estimation) were performed using a common reference, with all samples (including *Megaloceros* and modern cervids) mapped to the *Dama dama* assembly generated in this study. In contrast, analyses sensitive to reference divergence, including estimates of genetic diversity, inbreeding, and demographic history^28^, were performed using species-specific reference genomes. For these analyses, genomic data from eight modern cervid species with high-coverage (>20×) data were mapped to their respective conspecific assemblies.

Because no conspecific reference genome is available for *Megaloceros*, single representative individuals from each modern cervid species were also mapped to the *Cervus elaphus* reference genome to provide a comparative framework for evaluating reference-dependent effects. An overview of all reference–sample combinations and mapping results is provided in Supplementary Table S2. Although available, *Muntiacus crinifrons*, *M. muntjak*, and *M. vaginalis* were not mapped to conspecific assemblies due to their unusually large and highly rearranged chromosomes ^95^.

#### Mitochondrial genomes

Adapter sequences and low-quality bases (< Q30) were trimmed with cutadapt 1.18 ^67^. Untrimmed reads or reads <30bp in length were discarded. Trimmed reads were subsequently mapped to the *Megaloceros* mitochondrial genome (NCBI GenBank Accession: MW802558.1) using BWA v0.7.17 aln with default mapping parameters. Reads with a mapping quality below 30 were removed using Samtools v1.15.1. Duplicate reads (reads with the same start and end coordinates) were identified using the java program MarkDuplicatesByStartEnd.jar (https://github.com/dariober/Java-cafe/tree/master/MarkDupsByStartEnd) and removed. Mitochondrial consensus sequences were called using Samtools v1.15.1 with 85% majority rule for base calling and minimum coverage of 3x.

### Systematic relationships

#### Nuclear genomic phylogeny and gene flow analyses

To perform phylogenomic analyses, we created fasta files from one representative individual per species mapped to the *Dama dama* assembly using a consensus base call approach (-dofasta 2) in ANGSD v0.935 ^96^ and specifying the following parameters -minq 20 -minmapq 20 -setmindepthind 10. From the fasta files we generated 50 kb sliding windows with a 1 Mb slide using bedtools v2.29.1 ^85^ and the makewindows function to generate the coordinates for the windows and SAMtools faidx to extract the sequence. This led to 2,405 window alignments that we took as loci for phylogenomic analysis.

We implemented two methods of phylogenomic tree inference, accounting for different sources of error in the data ^97^. The first was a summary coalescent approach, which assumes that error arises mainly from signal discordance caused by incomplete lineage sorting. Locus phylogenetic trees were inferred for each window alignment under maximum likelihood as implemented in IQ-TREE v2.3 ^98^, assuming a GTR+R6 substitution model ^99^. These locus trees were used as input for summary-coalescent species tree inference as implemented in ASTRAL v5.8 ^100^. Secondly, we concatenated locus alignments under the assumption that error arises primarily from the use of a finite number of sites. The concatenated alignment was used for maximum likelihood phylogenetic inference with the same modelling approach as for locus trees. We then used concordance factor estimates for describing the decisiveness of the data in locus trees and alignment sites in describing the tree inferred ^101^. We also extracted discordance factors to quantify how strongly gene trees (or individual sites) support the two alternative topological resolutions around each internal branch of the species tree, providing a measure of imbalance in the underlying phylogenetic signal.

Bayesian molecular dating was performed using the concatenated alignment and a fossil calibration at the root of the process, balancing the informativeness and the placement certainty in the prior ^102^. This corresponded to *Ligeromeryx praestans* ^103^ and was represented as a uniform prior on the root age ranging from 19.5 to 17.2 Ma ^104^, with a soft maximum bound including 2.5% of the prior density. Molecular dating was implemented using approximate likelihood computation in two steps: first inferring branch lengths together with the gradient and Hessian matrix at the maximum likelihood estimate, followed by approximate likelihood MCMC sampling ^105^. This was implemented in MCMCtree as part of PAML v4.9 ^106^, using a birth-death branching process prior, a gamma-distributed uncorrelated branch rate prior, and a GTR+G6 substitution model. The MCMC chain was run for 50M steps sampling every 5K after a burn-in phase of 5M steps. Convergence to stationarity was confirmed by comparing two independent analyses and analysis of effective sample sizes and trace plots across inferred parameters.

We explored signals of gene flow between our selected species using Dsuite v0.4 r43^107^. Due to the large number of possible comparisons, and difficulties disentangling false positives that may arise due to ancient gene flow events, we performed the *f*-branch test ^107,108^. The test takes correlated allele sharing into account when visualising ƒ4-ratio results. As input, we created a multi-individual VCF file from the individuals mapped to the *Dama* assembly, containing only variant positions using freebayes v1.3.6^109^. We filtered to only include autosomal chromosomes, minimum mapping and base quality filters set to 20 (-q 20 - Q 20), only call heterozygous when the alternative allele frequency for a given site is >0.25 (--min-alternate-fraction 0.25), and remove all sites with any missing data. We ran the multi-individual VCF through Dtrios in Dsuite and specified the species tree calculated above as the most common topology and otherwise default parameters. We enabled the --ABBAclustering option to assess whether significant D-statistics could reflect mutational or reference biases rather than genuine introgression. We ran the output from Dtrios through *f*-branch and visualised the output using the dtools.py script from Dsuite.

#### Time-calibrated mitochondrial Bayesian analysis

The five newly reconstructed mitochondrial genomes from Rhine fossils were combined with 39 previously published *Megaloceros* mitochondrial sequences covering the species’ late Pleistocene Eurasian distribution ^20,26^. This dataset included four sequences from east Asian *Sinomegaceros* as well as mitochondrial genomes from four Irish individuals for which nuclear data are presented here and which were described previously. Two additional *Sinomegaceros* sequences (NCBI GenBank Accession: OR263894.1, OR263896.1) were excluded due to an excess of private SNPs. Ancient mitochondrial genomes were aligned using MAFFT v7.31071 ^110^. Poorly-aligning parts of the D-loop were removed from the alignment resulting in a total alignment length of 16,354 bp. The optimal substitution model under the Bayesian information criterion was determined using ModelFinder ^99^ as implemented in IQTREE 2.0.3 ^98^.

A time-calibrated Bayesian analysis was performed in BEAST 2.7.2 ^111^. Samples with reliable radiocarbon dates were used as tip calibrations to inform the evolutionary rate, whereas samples dated beyond the radiocarbon limit or lacking direct dates were assigned broad uniform priors (0–120 ka). For these latter samples, posterior median ages and 95% HPD intervals were estimated from the BEAST output. Posterior median ages inferred for undated or poorly dated specimens were 43.8 kya (95% HPD: 24.8–66.2 kya; CADG532), 41.4 kya (23.2–64.6 kya; CADG1199), 55.7 kya (41.0–70.9 kya; Ari42), 48.3 kya (32.6–64.3 kya; Ari50), 51.2 kya (37.8–64.0 kya; Ari5), and 97.8 kya (76.0–124.4 kya; Ari66). A Coalescence Bayesian Skyline tree model with a strict clock was applied and the clock rate was set to 1.65*10^-8^ substitutions/site/year ^26^. The MCMC chain was run for 100 million generations. Convergence and adequate sampling (ESS > 200) of all parameters were verified in Tracer 1.5.0.52 ^112^. The first 20% of trees were removed as burn-in, and the maximum clade credibility tree obtained from the posterior sample, with node heights scaled to the median of the posterior sample, using TreeAnnotator v2.7.1.

#### Population genomics

We evaluated the population relationships between our *Megaloceros* individuals by performing Principal Component Analyses (PCA). We computed two PCA, using either pseudohaploid base calls or genotype likelihoods computed with ANGSDv0.935 ^96^. We computed genotype likelihoods in ANGSD for all individuals specifying the parameters: minimum mapping and base qualities of 20 (-minmapQ 20 -minQ 20), calculate genotype likelihoods using the GATK algorithm (-GL 2), output a beagle genotype likelihood file (-doGlf 2), calculate major and minor alleles based on genotype likelihoods (-doMajorMinor 1), remove transitions (-rmtrans 1), only include SNPs with a p-values <1e-6 (-SNP_pval 1e-6), only consider autosomal chromosomes (-rf), a minimum minor allele frequency of 0.05 (-minmaf 0.05), skip triallelic sites (-skiptriallelic 1), only consider reads mapping to one region uniquely (-uniqueonly 1), and only consider sites if at least five individuals have coverage (-minind 5). The pseudohaploid base call was computed using the same parameters with the addition of -doIBS 2, -doCov 1, and only included SNPs if at least two individuals had the minor allele (-minminor 2). To construct covariance matrices from the genotype likelihoods datasets we used PCAngsd v0.98 ^113^. We evaluated the reliability of each PC axis using a Tracy-Widom test in R.

We further investigated population structure by computing a neighbour joining tree using the pseudohaploid base call (-doIBS 2) mentioned above and converted these into a distance matrix using the -makematrix 1 parameter in ANGSD. We converted the distance matrix into a neighbour joining phylogenetic tree using fastME v2.1.6.1 ^114^ and default parameters.

To quantify the level of divergence between populations, we used a consensus haploid base call in ANGSD and calculated the fixation index (*F*_ST_). To compute this, we specified the same parameters in ANGSD as used for the pseudohaploid PCA with the addition of outputting the consensus haploid base (-dohaplocall 2). We ran the resultant haploid output through the popgenWindows.py python script https://github.com/simonhmartin/genomics_general - downloaded June 2022), while specifying the population (Irish or German) for each individual, 100kb non-overlapping sliding windows in which to calculate *F*_ST_. Only windows with >100 variant sites were considered. Finally, we estimated the genetic diversity in each population via nucleotide diversity as the popgenWindows.py python script, used for the *F*_ST_ calculation above, also outputs nucleotide diversity for each window.

#### Selection

We constructed a multispecies dataset for positive selection analyses including human (*Homo sapiens*; hg38), white-tailed deer (*Odocoileus virginianus*), Père David’s deer (*Elaphurus davidianus*), sika deer (*Cervus nippon*), Reeves’s muntjac (*Muntiacus reevesi*), roe deer (*Capreolus capreolus*), Eurasian elk (*Alces alces*), Eld’s deer (*Rucervus eldii*), red deer (*Cervus elaphus*), reindeer (*Rangifer tarandus*), hog deer (*Axis porcinus*), fallow deer (*Dama dama*), Chinese water deer (*Hydropotes inermis*), and *Megaloceros giganteus* (the consensus sequence constructed above for the phylogenomic analyses). We generated whole-genome alignments to the human reference using Segalign^115^ and inferred orthologous gene projections with TOGA v1.1.7^116^. A total of 19,465 annotated human genes were provided as input to TOGA for each species. From the resulting predictions, we retained only genes classified as “one-to-one” orthologs. The final set of 11,906 orthologous genes represents the intersection of these one-to-one orthologs across all analysed species. We extracted codon-aligned orthologous sequences using the TOGA *extract_codon_alignment.py* script, aligned them with MACSE v2^117^, and filtered the alignments using Gblocks v0.91b^118^.

We assessed lineage-specific positive selection in *Megaloceros* using two complementary codon-based approaches implemented in HyPhy v2.5.48^119^. First, we evaluated branch-specific rate variation under the MG94 codon model using the *fitMG94* module. For each gene, we fitted a one-ratio model assuming a single dN/dS ratio (ω) across the phylogeny to estimate tree-wide selective pressure, and we compared it to a free-ratio model allowing branch-specific ω values. We considered genes showing ω > 1 on the *Megaloceros* branch and a significantly better fit of the free-ratio model relative to the one-ratio model, as assessed by a likelihood ratio test (LRT), as candidates for lineage-specific positive selection.

Second, we tested for episodic positive selection acting on the *Megaloceros* lineage using the Branch-site Unrestricted Statistical Test for Episodic Diversification (BUSTED), designating *Megaloceros* as the foreground branch. We inferred genes with BUSTED *P* `≤ 0.05 to have experienced episodic positive selection. We further identified positively selected sites within candidate genes using the MEME method`^120^. We considered genes supported by both elevated branch-specific ω values and significant BUSTED results to be the most robust candidates for positive selection.

Using Metascape^29^, the genes identified as undergoing episodic positive selection were subjected to functional enrichment analysis. Significantly enriched terms and pathways (q < 0.05) were identified across Gene Ontology biological processes, as well as KEGG, Reactome, and WikiPathways categories.

#### Genetic diversity and inbreeding

We jointly calculated genome-wide heterozygosity and runs of homozygosity (ROH) for individuals mapped to their conspecific assembly using a modified version of a previously published pipeline that utilises allele counts^45^. Species we analysed included: *Axis porcinus* (Hog deer), *Capreolus pygargus* (Eastern roe deer), *Cervus canadensis* (wapiti), *Cervus elaphus* (red deer), *Cervus hanglu yarkandensis* (Yarkand deer), *Muntiacus gongshanensis* (Gongshan muntjac), *Muntiacus reevesi* (Reeves’ muntjac), *Rangifer tarandus platyrhynchus* (Svalbard reindeer), *Dama dama* (European fallow deer), *Megaloceros giganteus* (giant deer). In brief, we used ANGSD to directly perform allele counts on the autosomes of the mapped bam files using the following command and filters: -minQ 20 -minMapQ 20 - uniqueOnly 1 -remove_bads 1 -doCounts 1 -dumpCounts 4. We set the minimum depth as half the mean coverage of the given individuals. We only called heterozygous sites if the minor allele was >5%. We visualised all resultant minor allele frequencies for heterozygous base calls (Supplementary Fig S7) and chose a 25% minor allele frequency threshold to call a site as heterozygous. We reran the analysis with a 25% threshold to call a site as heterozygous. A site with an allele found at a frequency of >75% was called homozygous. Any sites that did not have two alleles with a frequency of >25% each or one allele with >75% frequency was counted but not given a base call. We divided the total number of heterozygous bases by the total number of sites that passed the initial ANGSD filtering in the individual. From the base call output, we counted runs of homozygosity in 100 kb windows, only considering a window as ROH if the proportion of heterozygous sites in that window was <0.0001 and the window contained at least 25 kb of called sites. We further counted the number of consecutive 100 kb windows in ROH. To investigate the role of reference genome selection and aDNA damage, we reran this analysis using the data mapped to a heterospecific reference genome and only considering transversion heterozygous sites.

We also performed heterozygosity estimations on a modified version of the above allele count analysis on the low-coverage German individuals Meng14, Meng16 (5.64x and 4.56x respectively), and a downsampled version of MG3 which we downsampled to ∼5x with SAMtools to gain insights into the relative diversity levels between our two *Megaloceros* datasets. We modified the approach to only include sites with at least 5x coverage (-setMinDepthInd in ANGSD) and if any site was above 5x, then it was downsampled to 5x (-capdepth in ANGSD). We extracted each site with a heterozygous base call and investigated whether there were one or two alternative alleles. Due to vastly different levels of single alternative allele sites (Supplementary table S4), likely due to various factors including differences in wet laboratory processes and sequencing error rates, we decided to only count a site as heterozygous if two out of the five reads contained an alternative allele. We restricted this analysis to transversion only heterozygous calls to avoid differences in levels of aDNA damage.

As we lack a conspecific reference genome for *Megaloceros*, we used the data mapped to *Dama dama*. However, there is potential for reference bias in heterozygosity base calls when mapping to a distant reference genome ^28^. Therefore, we evaluated the impact of the reference genome by recalculating heterozygosity and ROH for all modern individuals mapped to both *Dama dama* and *Cervus elaphus*. We investigated whether distance to the reference genome gave predictably biased results by plotting relative difference to the ‘true’ heterozygosity obtained when mapping to a conspecific reference and the divergence time between the species of interest and the reference genome. As we did not find any clear association between these two variables, we recalculated the heterozygosity for the *Megaloceros* using the mean (1.15) and maximum (1.48) deviations from the ‘true’ heterozygosity obtained from the modern individuals. We recalculated ROH in the *Megaloceros* by increasing the minimum cutoff for a ROH window from <0.0001 to <0.000115 and <0.000148 respectively. To evaluate whether reference bias and its resultant higher levels of heterozygosity were interrupting long ROH, we visualised ROH for MG3, *Dama dama* mapped to itself, *Dama dama* mapped to *Cervus elaphus, Cervus canadensis* mapped to itself, and *Cervus canadensis* mapped to *Cervus elaphus.* Based on these results, we additionally implemented a one-window bridging rule, whereby a single non-ROH 100 kb window flanked on both sides by ROH windows was reclassified as ROH. This additional analysis was only performed on MG3, *Dama dama* mapped to itself, *Dama dama* mapped to *Cervus elaphus, Cervus canadensis* mapped to itself, and *Cervus canadensis* mapped to *Cervus elaphus*.

To complement allele-count–based estimates of genetic diversity in the lower-coverage Megaloceros individuals, we additionally inferred heterozygosity from genotype likelihoods using ATLAS (tasks *estimateErrors* and *summaryStats*), which explicitly accounts for sequencing error and post-mortem damage ^94,121^ (bitbucket.org/wegmannlab/atlas, commit 7b82b51b). In brief, this approach first estimates post-mortem damage patterns and sequencing error rates, which are used to calculate accurate genotype likelihoods. From these, heterozygosity is subsequently inferred under Felsenstein’s substitution model. Error rates were modeled using the empiric function of the following covariates: quality score, position in thread, mapping quality, fragment length and context (i.e. the previous base sequenced). The model was inferred from all sequence data on two 10Mb regions on chromosome 5. The heterozygosity was then inferred in 493 windows of 5Mb along all autosomes. To avoid mapping errors, we excluded all sequencing reads of which more than 5% of all mapped bases did not match the reference or for which more than 10% of all bases were soft clipped (options --filterSoftClips ",0.1" and --filterMappingMismatches ",0.05"). A negative binomial distribution was fitted over the combined depth of all ancient samples using the ATLAS tasks pileup and pileup2Bed. All sites falling outside the 99% quantile of the fitted distribution were masked.

To confirm the accuracy of the inferred post-mortem damage patterns and sequencing error rates, and consequently of diversity estimates, heterozygosity was also inferred from sequencing data down-sampled to depths 50,20,10,5,2,1,0.5,0.2,0.1 using the task summaryStats with the option --depth. Under accurate estimates of post-mortem damage patterns and sequencing error rates, the diversity estimates are expected to be rather insensitive to the sequencing depth, as was confirmed for the samples used here (Supplementary Fig. S12).

### Genetic load

#### Temporal dynamics of genetic load in *Megaloceros* from Germany and Ireland

To investigate temporal dynamics of genetic load in *Megaloceros*, we analysed a subset comprising four German individuals (36.7–46.6 kya BP; 3.6×–5.6× coverage), three low-coverage Irish individuals (11.0–11.8 kya BP; 0.4×–5.6× coverage), and one high-coverage Irish individual (11.1 kya BP; 48× coverage), together with modern *D. dama* and *C. elaphus* genomes used as outgroups. We generated a joint VCF from all individuals mapped to the *Dama dama* assembly using FreeBayes v1.3.6. Variant calling was restricted to autosomal regions and applied minimum mapping and base quality thresholds of 20 (--min-mapping-quality 20; --min-base-quality 20). To reduce false heterozygous calls arising from sequencing error and ancient DNA damage, we required a minimum alternative allele fraction of 0.25 (--min-alternate-fraction 0.25). We excluded all transition polymorphisms using BCFtoolsv1.16 ^122^ and retained only transversion sites for downstream analyses. We further filtered the dataset to retain only sites present in all individuals (no missing data) and removed variants with a maximum mean coverage across samples exceeding 70× using VCFtools ^123^. The script vcfallelicprimitives in the vcflib package ^124^ was used to split the multiple nucleotides variants in single nucleotide variants, which were filtered for overall Phred-quality > 30, strand bias (SAF > 0 & SAR > 0) and read position bias (RPL > 0 & RPR > 0) using the script vcffilter in the vcflib package. Only variants identified as SNPs were retained. The dataset was further filtered to include only biallelic SNPs using vcftools and the functional effect of the identified genetic variants was predicted using SnpEff ^125^. SnpEff classifies variants into four categories of putative impact on fitness: high, moderate, low, modifier. High-impact variants are expected to have severe effects on protein function (e.g., loss-of-function) while moderate-impact variants typically correspond to non-synonymous changes with smaller functional effects. Low-impact variants include synonymous substitutions and modifier variants generally fall outside genic regions and are assumed to be largely neutral. If a variant fell into multiple impact categories, we retained the first category identified according to SnpEff’s hierarchy. Concerning high and moderate impact variants, the derived state is commonly assumed to be the deleterious one. We considered the allele in *D. dama* and *C. elaphus* as the ancestral state only if fixed in both. Ancestral-derived state polarization and genotype counts summaries were performed using GenoLoader (v3.2, https://github.com/emitruc/genoloader).

As raw counts of homozygous genotypes were strongly correlated with individual sequencing depth (Supplementary Fig S13), we performed a random pseudo-haploidization of the individual genotypes (a common approach when analyzing low coverage ancient data, see e.g. ^126^) and counted the ancestral and derived allele per fitness impact class. For each variant site, we randomly sampled one read at each individual genotype and checked whether it was supporting the ancestral or derived allele using the option low_cov in the write_gt GenoLoader function. Using pseudo-haploid data, we calculated the R_xy_ statistic ^127^ for each fitness impact class, setting the Irish sample as x population and the German samples as the y population. Then, we calculated the normalized statistic R’_xy_, where R_xy_ values for low, moderate and high impact categories were normalized using the corresponding value obtained for the modifier class, which are assumed to represent neutral variation. To account for stochastic variation introduced by allele resampling, the pseudo-haploidization procedure was repeated 100 times. At each iteration R_xy_ and R’_xy_ were calculated for all fitness impact classes. This approach allowed us to compare relative levels of derived variation between temporally distinct populations of *Megaloceros,* while accounting for the bias introduced by the uneven sequencing coverage in the samples, and to evaluate whether our data shows evidence of genetic load accumulation or depletion after the colonization of Ireland from mainland Europe and while approaching the extinction of the Irish population.

#### Analysis of genetic load in *Megaloceros* compared to extant cervid lineages

To evaluate genetic load in *Megaloceros* relative to other cervid species with varying demographic histories and extinction risks, we analysed a dataset comprising the high-coverage Irish *Megaloceros* genome together with one >20× genome per species from a panel of extant cervids (*Axis porcinus, Dama dama, Cervus canadensis, C. elaphus, C. hanglu, Muntiacus crinifrons, M. gongshanensis, M. muntjak, M. reevesi,* and *M. vaginalis*). All genomes included had average coverage between 20× and 48× (mean: 26×). We additionally included *Bos taurus, Capreolus capreolus,* and *Rangifer tarandus* as outgroups for ancestral state inference.

Variants were called jointly from individuals mapped to the *Dama dama* assembly using the same FreeBayes-based pipeline and quality filters described above but with the inclusion of transitions. After restricting the dataset to this subset of genomes, we retained only autosomal, high-quality biallelic SNPs with no missing data and applied a minimum per-genotype coverage threshold of 7×.

To infer ancestral and derived states, we required sites to be invariant in the outgroup taxa, with the outgroup allele defined as ancestral. Sites were retained if polymorphic exclusively within one of two predefined ingroups: (i) *Dama, Cervus, Megaloceros,* and *Axis*; or (ii) all *Muntiacus* species, thereby restricting analyses to lineage-specific variation arising after the split between these clades. Ancestral–derived polarization, variant filtering, and genotype count summaries were performed using GenoLoader.

We approximated the genetic load by i) counting the number of homozygous genotypes for the derived allele at high-impact sites in each lineage (raw counts), and ii) by calculating the ratio of homozygous genotypes for the derived allele at high-impact sites over homozygous genotypes for the derived allele at modifier-impact sites (normalized load). Because mutations at modifier sites are expected to represent mostly neutral variation, negative deviations from neutrality in this ratio may indicate relaxed purifying selection or evidence of purging. To estimate the significance of the pattern across the genome, we performed a bootstrap designed as follows: we randomly sampled 1,000 sites per ingroup for the high-impact category; then we randomly sampled two sets of 1,000 modifier-impact sites to be used as the neutral (null) distribution. This process was repeated 100 times. To ensure that our sampled modifier sites were representative of the broader modifier class, we additionally calculated a control ratio between the two independent sets of 1,000 modifier sites. The latter were consistently close to 1, confirming the stability of our neutral baseline set by the modifier variants (Supplementary Fig S16).

#### Demographic history

We inferred the long-term demographic history of our *Megaloceros* MG3 genome using a pairwise sequential markovian coalescent model PSMC ^128^. We called diploid genome sequences using SAMtools and BCFtools, specifying a minimum quality score of 20, minimum coverage of 10, and specified only to autosomes >100kb in length. We subsequently filtered the vcf file utilising the output of the heterozygosity base call pipeline from above to only include heterozygous sites that had a minor/alternative allele frequency >25%. We ran PSMC specifying standard atomic intervals (4+25*2+4+6), a maximum number of iterations of 25 (-N), maximum 2N0 coalescent time of 15 (-t), an initial theta/rho ratio of 5 (-r), and performed 50 bootstrap replicates. We checked the PSMC results for overfitting, ensuring that after 20 rounds of iterations, at least 10 recombination events were inferred within the intervals spanned by each parameter. As no mutation rate for *Megaloceros* was available, we calculated one from the divergence time between *Dama* and *Megaloceros* estimated in this study and their autosome-wide pairwise difference (PWD). We calculated the PWD using a consensus base call approach on the autosomal chromosomes in ANGSD (-doIBS 2) and the following parameters: -minmapQ 20 -minQ 20 -doCounts 1 -GL 2 - doMajorMinor 1 -docov 1 -makematrix 1 -uniqueonly 1 -minind 2. We recovered a PWD=of 0.0141275 and using the 3.53Ma divergence and the equation PWD/2/divergence time we got a mutation rate per site per year of 2e-9. As *Megalcoceros* is estimated to have been around 450–600 kg, and comparable in size to large moose ^8^, we estimated a generation time of 10 years based on the generation times for *Alces americanus* (mean weight 541.5 kg), and *Alces alces* (mean weight 461.9 kg) of 9.34 and 10.20 respectively ^129^. This gave us a mutation rate per site of 2e-8 per generation that we used to plot our results.

We additionally inferred demographic histories for four extant cervids selected for their relevance to *Megaloceros*: *Cervus elaphus* (red deer), with a broadly overlapping Eurasian distribution ^20,130^; *Capreolus pygargus* (eastern roe deer), associated with comparable forest–steppe habitats ^20,131^; *Cervus canadensis* (wapiti), an ecologically similar large-bodied mixed feeder ^132,133^; and *Dama dama* (fallow deer), the closest living relative of *Megaloceros*. We generated the PSMC outputs using the same approach as for *Megaloceros*. We plotted the red deer with a generation time of 6.3 years ^134^ and therefore mutation rate of 1.26e-8 per generation, wapiti with the same generation time and mutation rate as the red deer, eastern roe deer with a generation 6 years ^135^ and therefore mutation rate of 1.2e-8 per generation, and the fallow deer with a generation time of 9 years ^129^ and therefore mutation rate of 1.8e-8. We used the same mutation rate per year as for *Megaloceros*.

## Data Availability

Raw sequencing reads will be available upon acceptance.

## Acknowledgements

We thank Jes Rasmussen for allowing us access to the *Dama dama* individual used for the de novo assembly. We thank Matthew Teasdale for assistance with sample screening. We thank Ronny Friedrich and Susanne Lindauer from the Curt-Engelhorn-Zentrum Archäometrie gGmbH, Mannheim, Germany, for assistance with the radiocarbon dating of the German *Megaloceros* individuals.

## Author contributions

Conceptualization: M.V.W., M.T.P.G., M.-H.S.S., D.G.B., M.H.

Formal analysis: M.G., Z.L., E.T., P.A., S.K., D.Du., D.W., A.F., G.G., K.M.P., B.C., R.K., M.V.W.

Investigation: K.G.D., V.M., F.A., I.K., J.N., P.P., S.S.T.M.

Resources: N.T.M., W.R., D.Do.

Writing – original draft: M.V.W.

Writing – review & editing: All authors

Supervision: E.T., L.L., W.W., E.D.L., B.S., D.G.B., M.H., M.T.P.G., M.V.W.

