## Supplementary information for "Evolutionary history and genomic vulnerability of the extinct giant deer *Megaloceros giganteus*"

**Supplementary Figures**

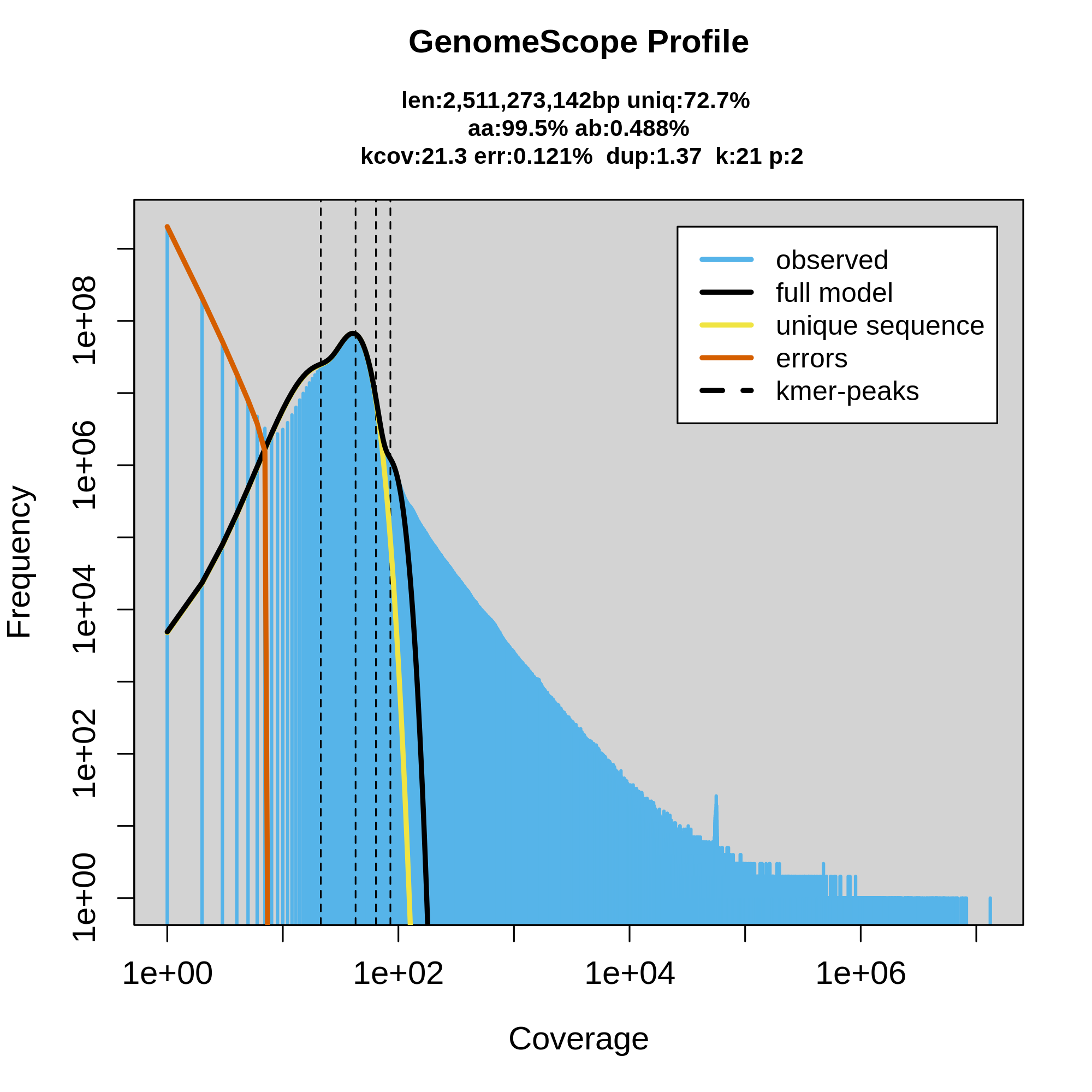

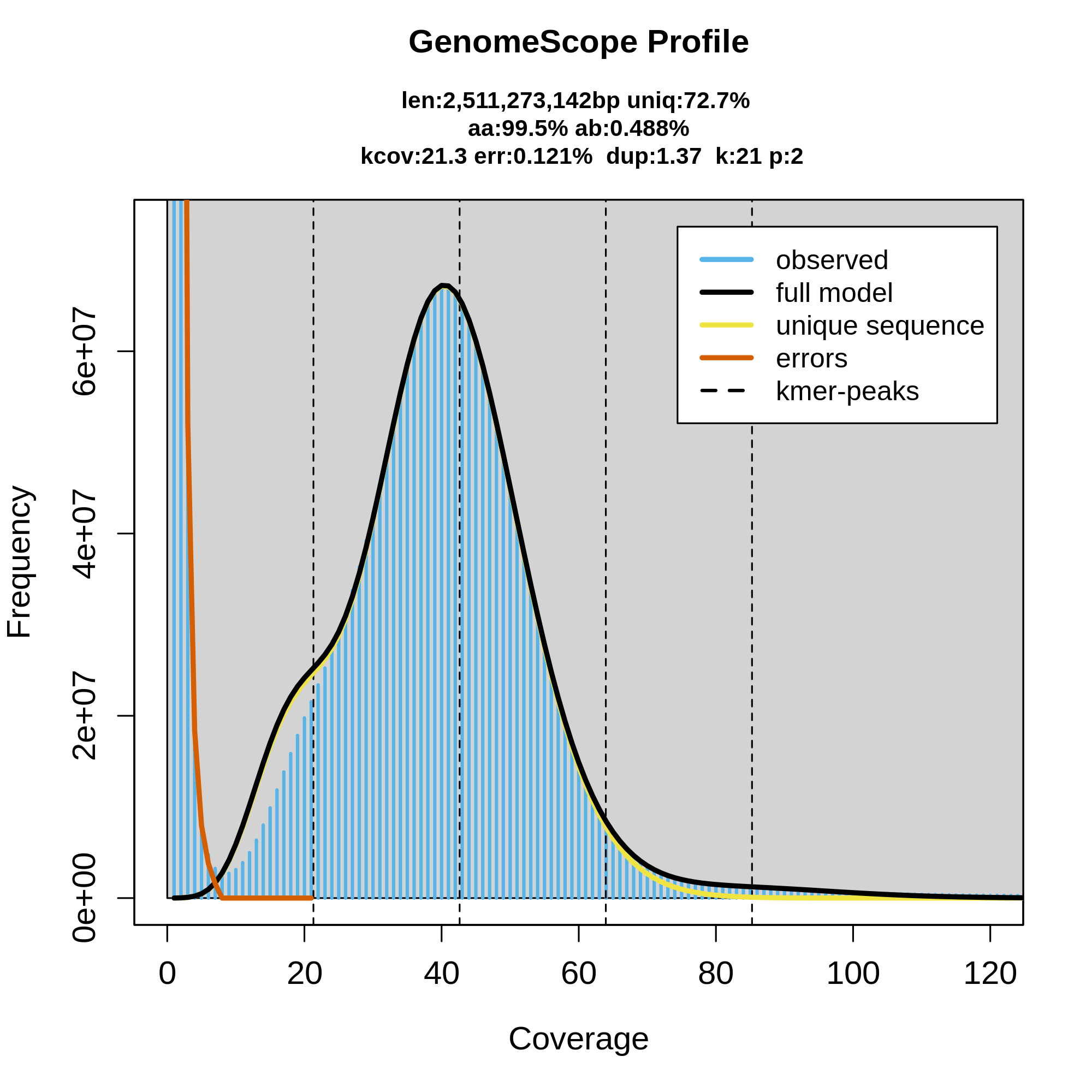

**Supplementary Fig S1.** **GenomeScope profile of 23-mer frequencies derived from PacBio HiFi reads of the *Dama dama* reference individual.** The figure shows the observed k-mer distribution (blue), the fitted GenomeScope model (black), and inferred components including unique sequence and sequencing error contributions. The lower panel provides a zoomed view of the main coverage peak.

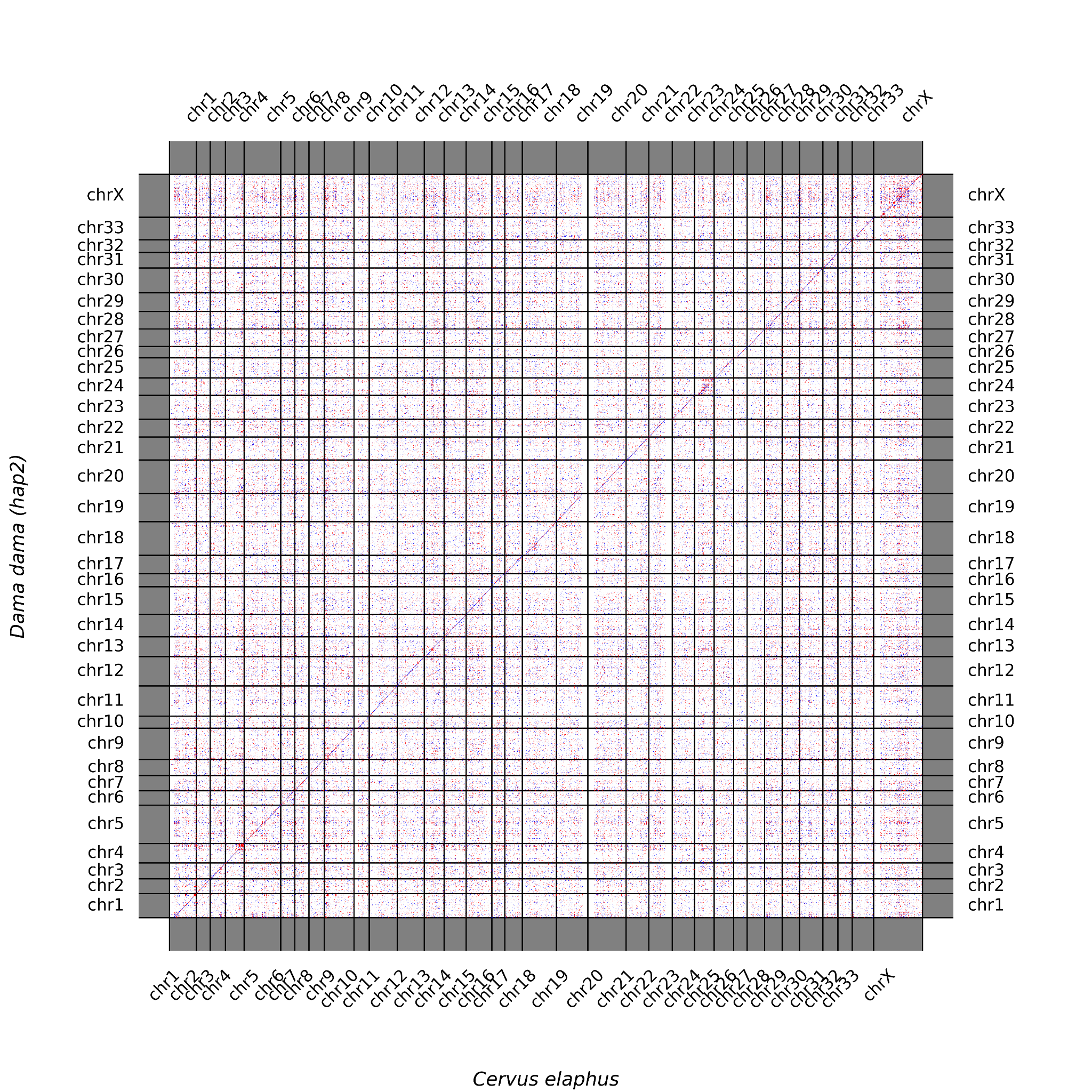

**Supplementary Fig S2. Hi-C–based synteny between the maternal pseudo-haplotype assembly of *Dama dama* and the reference assembly of *Cervus elaphus* (GCA_910594005.1), illustrating conserved chromosomal structure and correspondence between assemblies.**

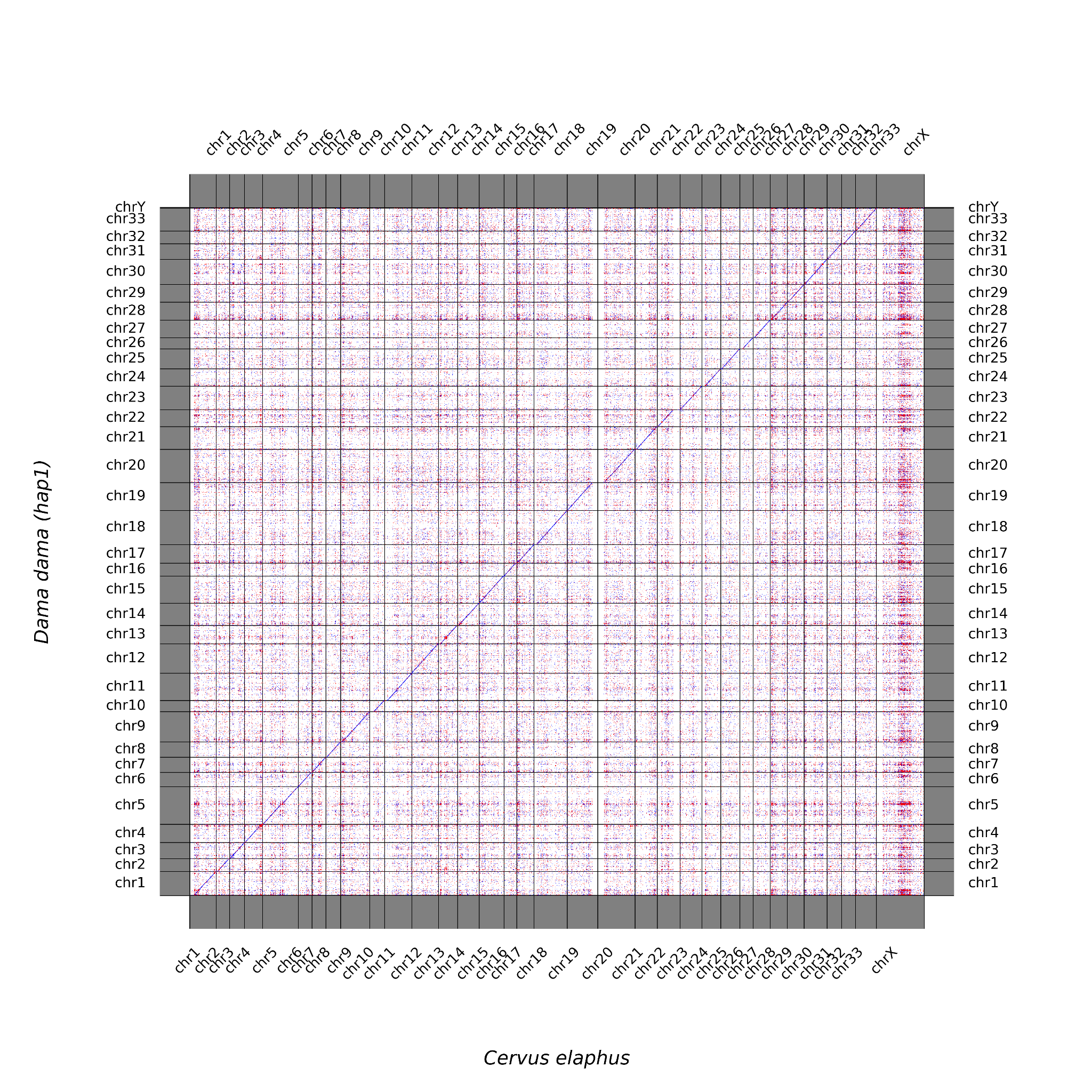

**Supplementary Fig S3.** **Hi-C–based synteny between the paternal pseudo-haplotype assembly of *Dama dama* and the reference assembly of *Cervus elaphus* (GCA_910594005.1), illustrating conserved chromosomal structure and correspondence between assemblies.**

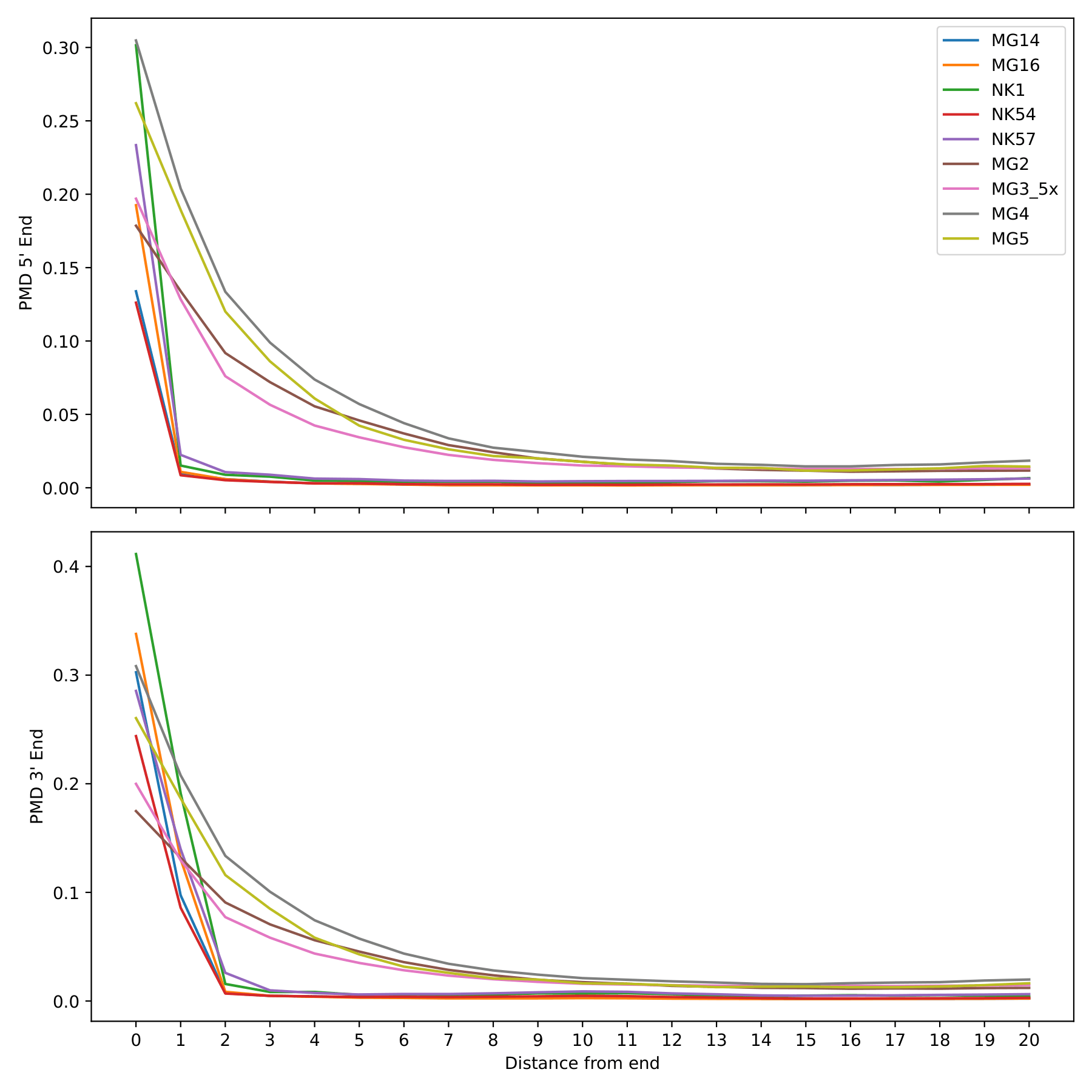

**Supplementary Fig S4. Postmortem damage (PMD) profiles for all nine *Megaloceros* individuals.** Lines represent nucleotide misincorporation patterns along read positions. MG3 was downsampled to ~5× coverage. The upper and lower panels show damage patterns at the 5′ and 3′ ends of reads, respectively.

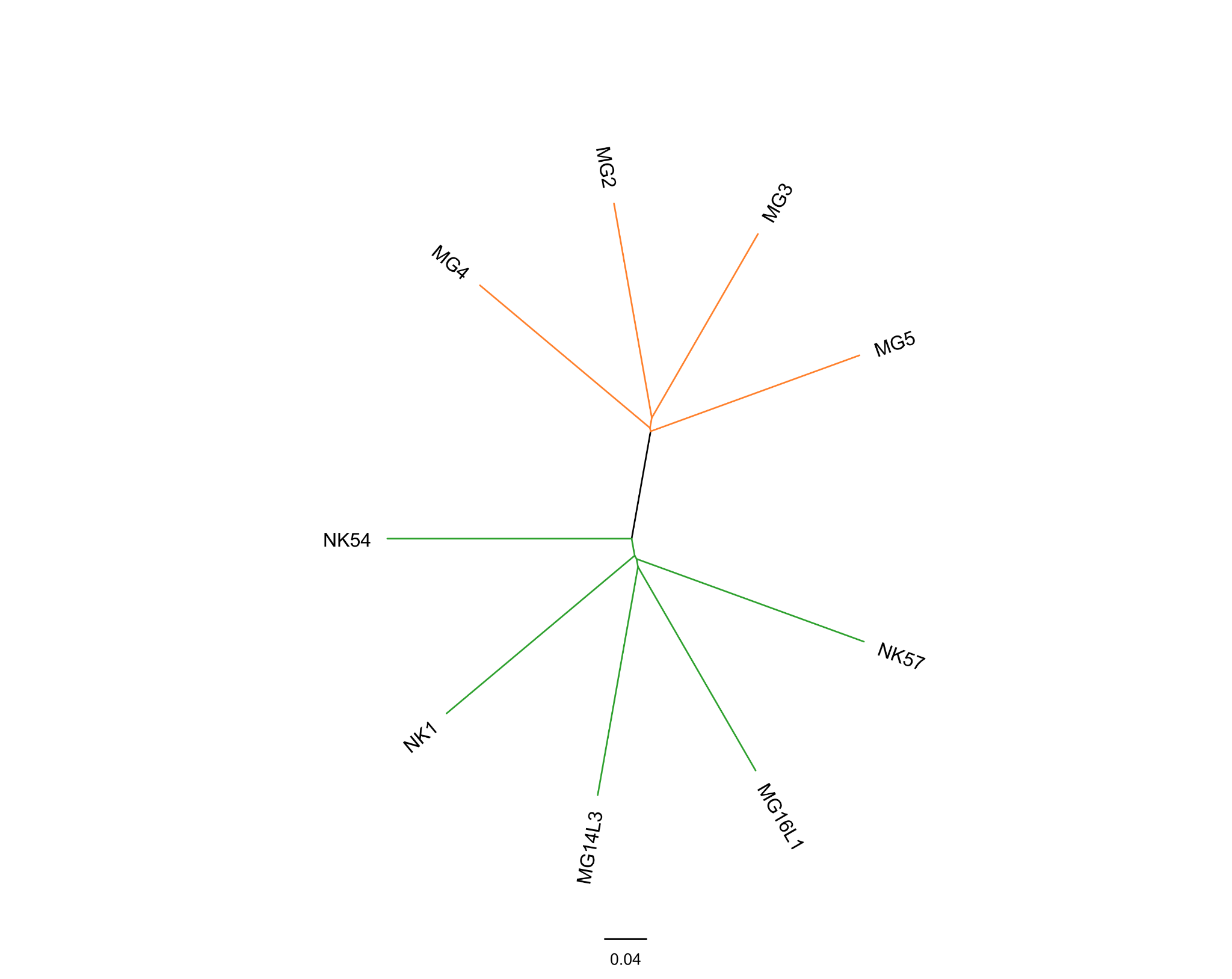

**Supplementary Fig S5.** **Unrooted neighbour joining tree of all *Megaloceros* individuals included in this study, based on nuclear genomic pseudohaploid base calls at sites where the minor allele is found in at least two individuals**. Irish individuals are indicated in orange, German individuals in green.

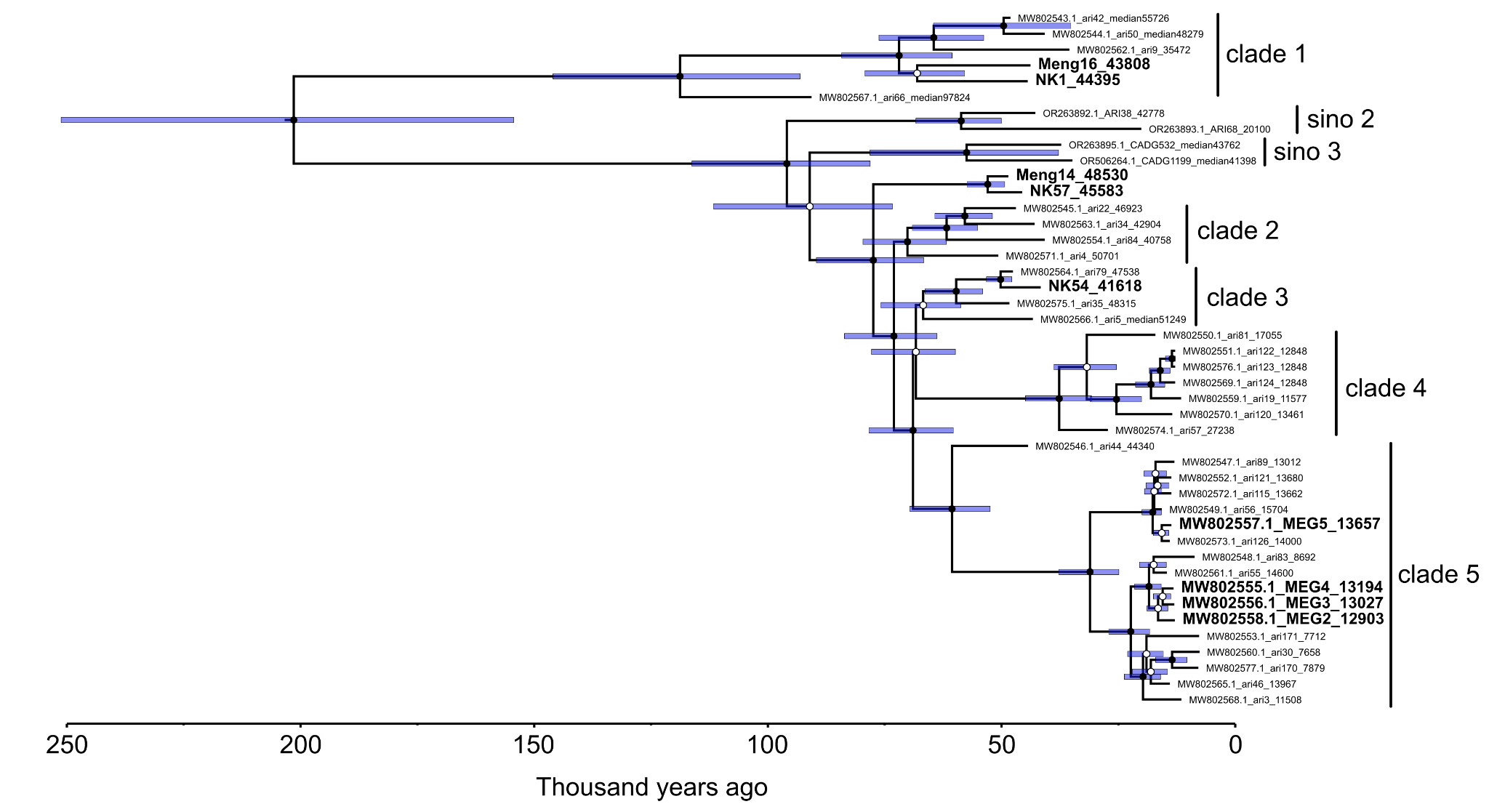

**Supplementary Fig S6.** Time-calibrated mitochondrial Bayesian phylogeny. Samples with nuclear data are in bold. Clade definitions follow Rey-Iglesia et al. (5; clade 1-5) and Xiao et al. (12, sino 2-3). Closed circles indicate node support with posterior probability >0.99, open circles indicate node support with posterior probability <0.99. Blue bars represent highest posterior density of node dates.

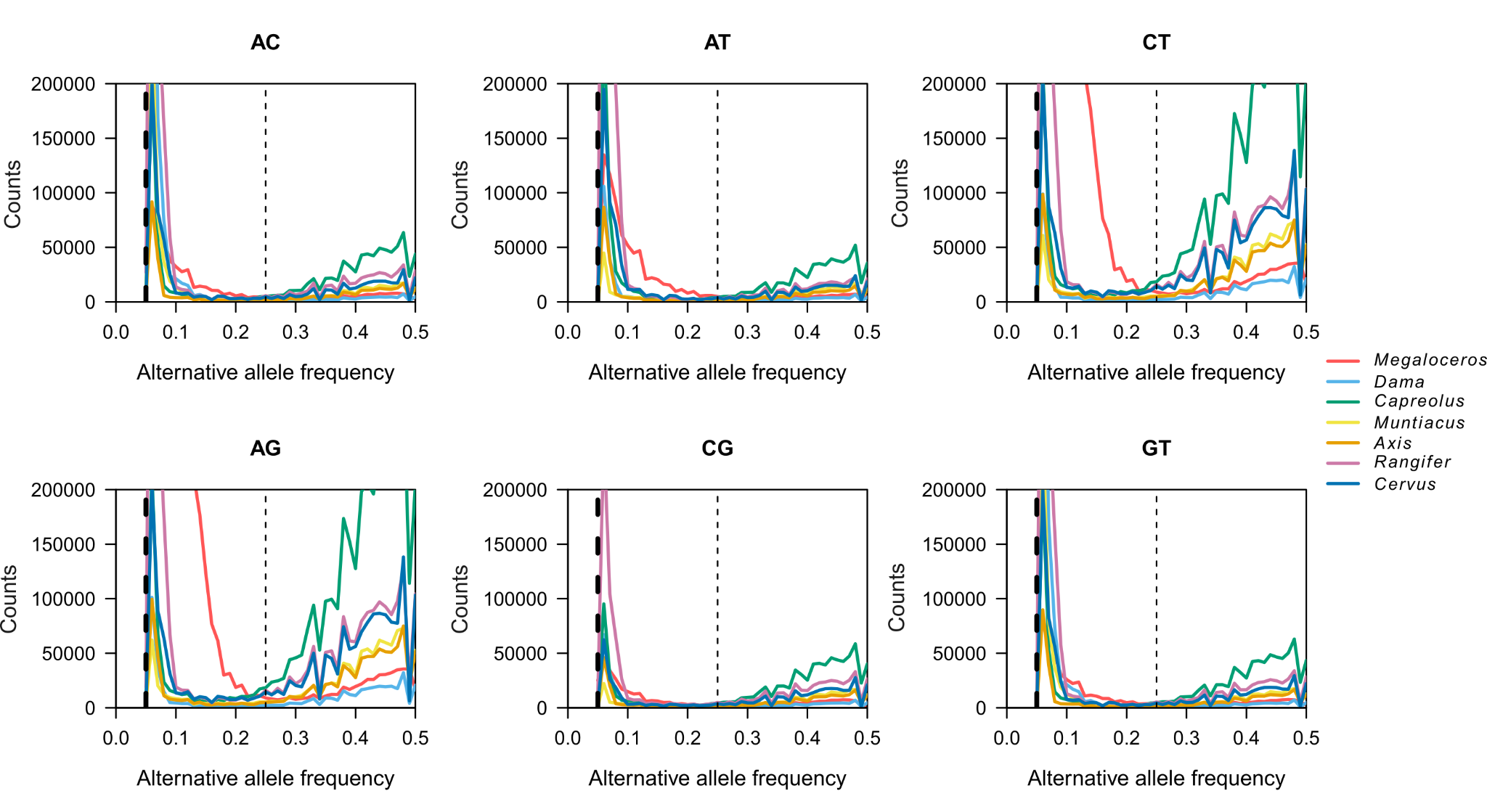

**Supplementary Figure S7:** **Counts of alternative alleles separated based on their frequencies for a single individual per genus.** *Cervus* shows *C. canadensis.* *Muntiacus* shows *M. gongshanensis*. The dotted line shows the 0.25 cutoff used in this study.

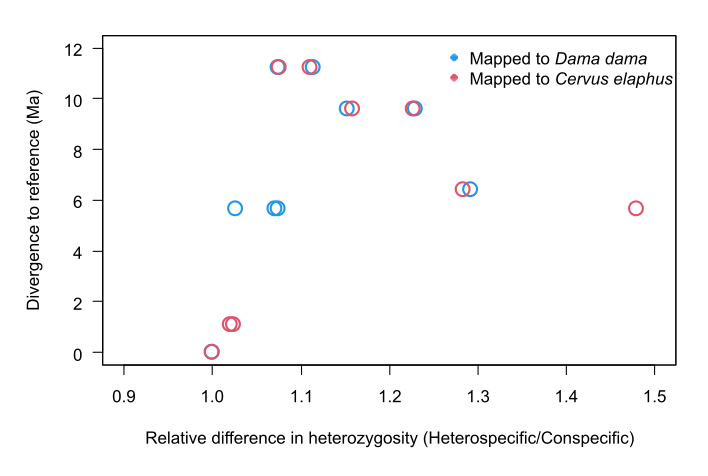

**Supplementary Fig S8: Comparisons between divergence to the reference genome relative to difference in heterozygosity.** Each point represents a modern cervid individual mapped to both a conspecific and a heterospecific (*D.dama* or *C.elaphus*) reference genome. The x-axis shows the ratio of autosomal heterozygosity estimated using the heterospecific reference relative to the conspecific reference (heterospecific / conspecific), reflecting the magnitude of reference bias. The y-axis shows the divergence time (Ma) between the focal species and the heterospecific reference genome.

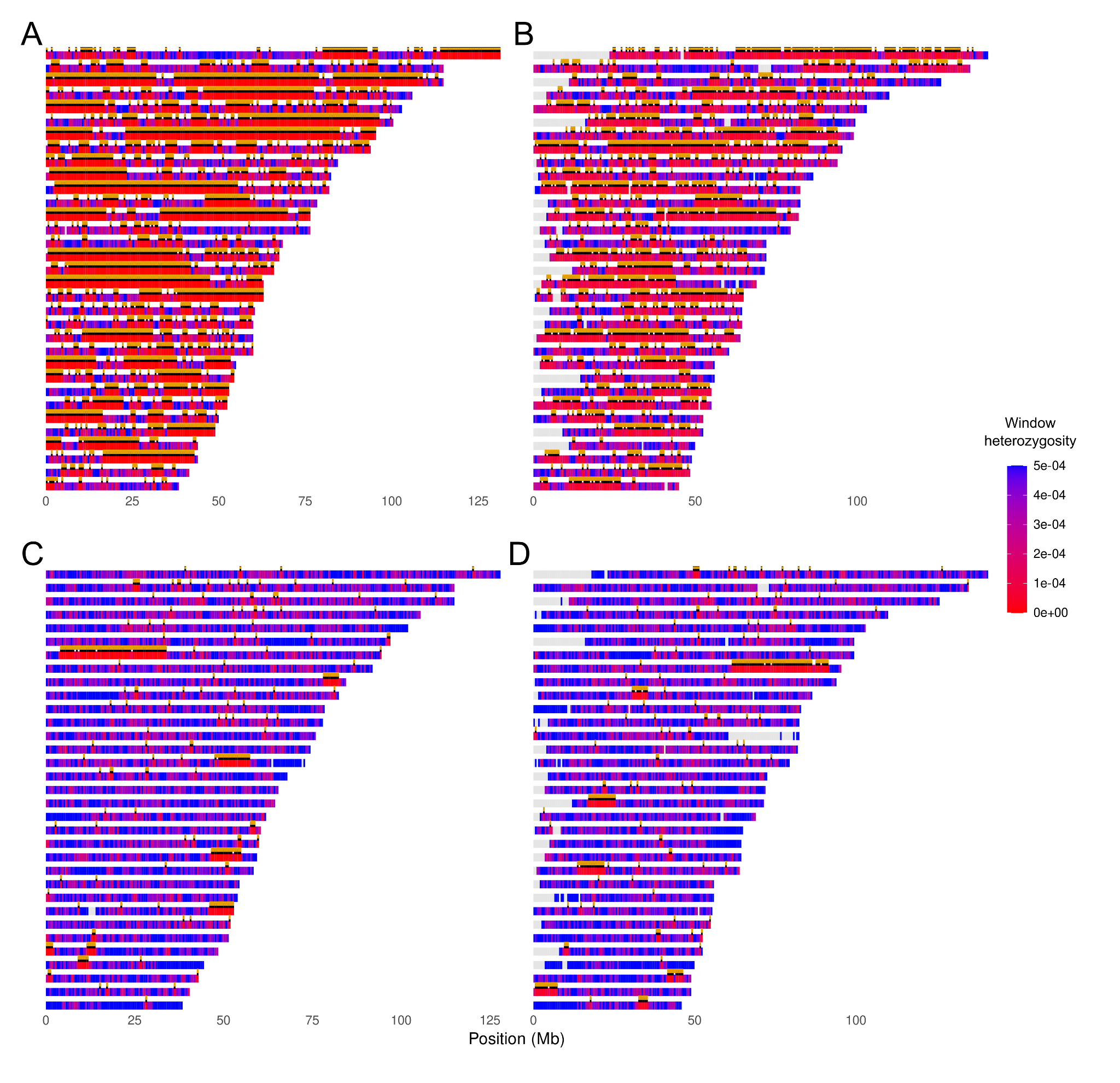

**Supplementary Figure S9: Visualisation of heterozygosity across the genomes of A)** *Dama dama* mapped to itself, **B)** *Dama dama* mapped to *Cervus elaphus*, **C)** *Cervus canadensis* mapped to itself, and **D)** *Cervus canadensis* mapped to *Cervus elaphus.* Any windows with a value >0.005 were given a value of 0.005 to ease visualisation.

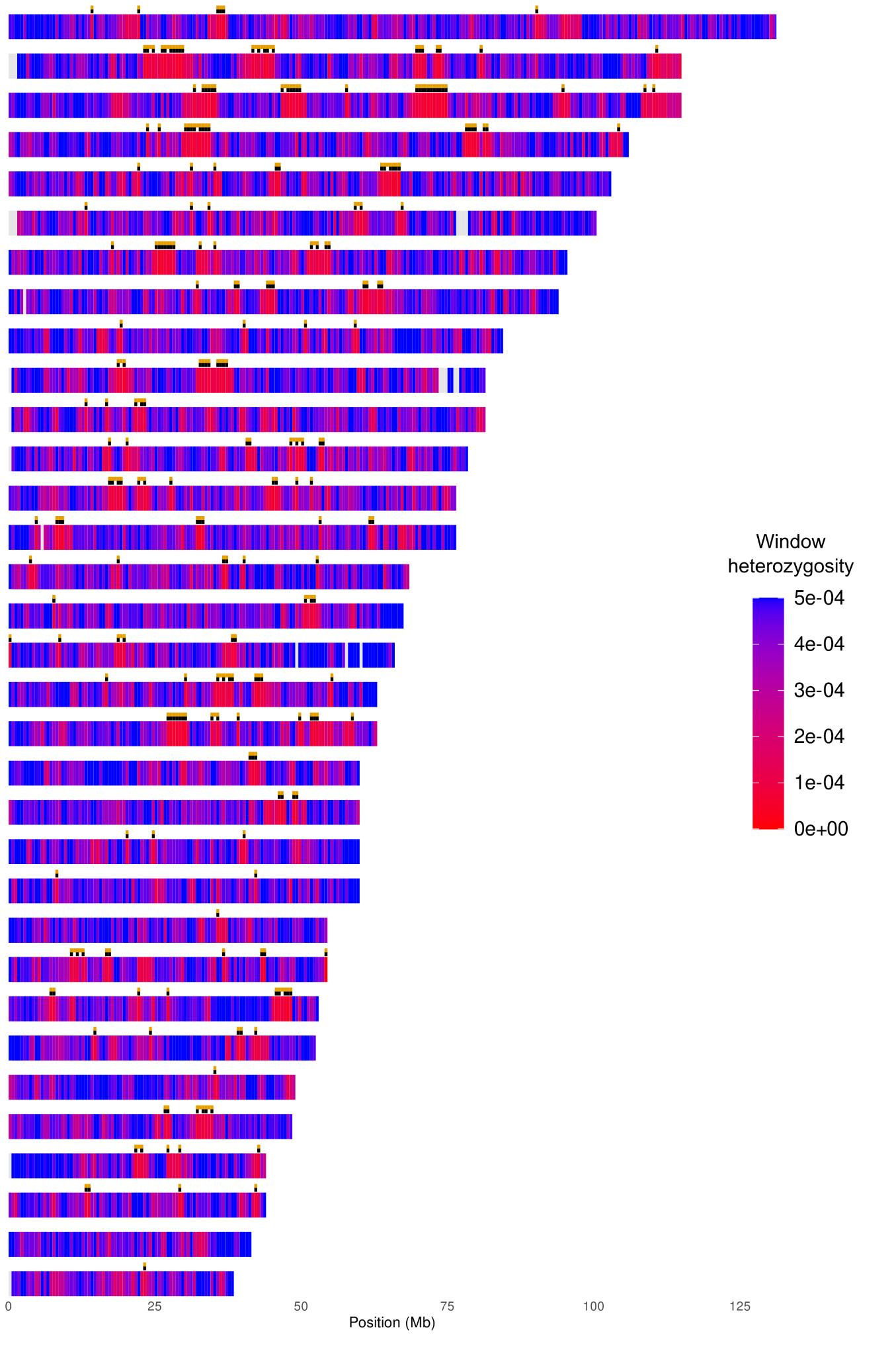

**Supplementary Figure S10:** Distribution of heterozygosity across the genome of high-coverage *Megaloceros* individual MG3. Any windows with a value >0.005 were given a value of 0.005 to ease visualisation and windows with missing data are depicted in white. Black ticks above chromosomes indicate windows classified as ROH (heterozygosity <= 0.0001), and orange ticks indicate ROH after one-window bridging (a single non-ROH window reclassified as ROH when flanked by ROH windows on both sides).

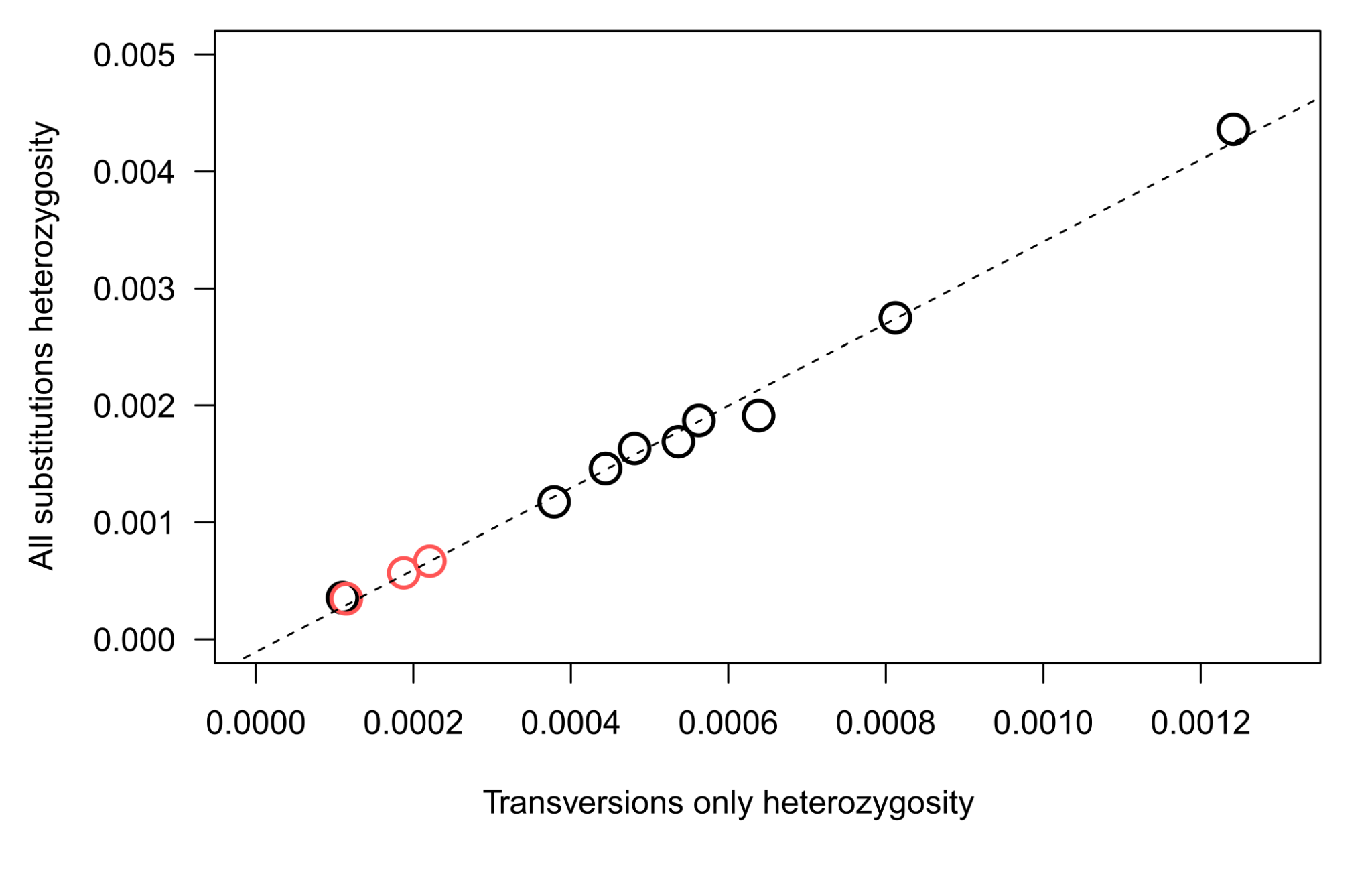

**Supplementary Fig S11: Comparison of heterozygosity estimates calculated using transversions only (x-axis) versus both transversions and transitions (y-axis) for nine cervid species (black open circles) and three *Megaloceros* replicates generated using different reference bias correction values (red open circles).** The dotted line indicates the linear regression across all data points.

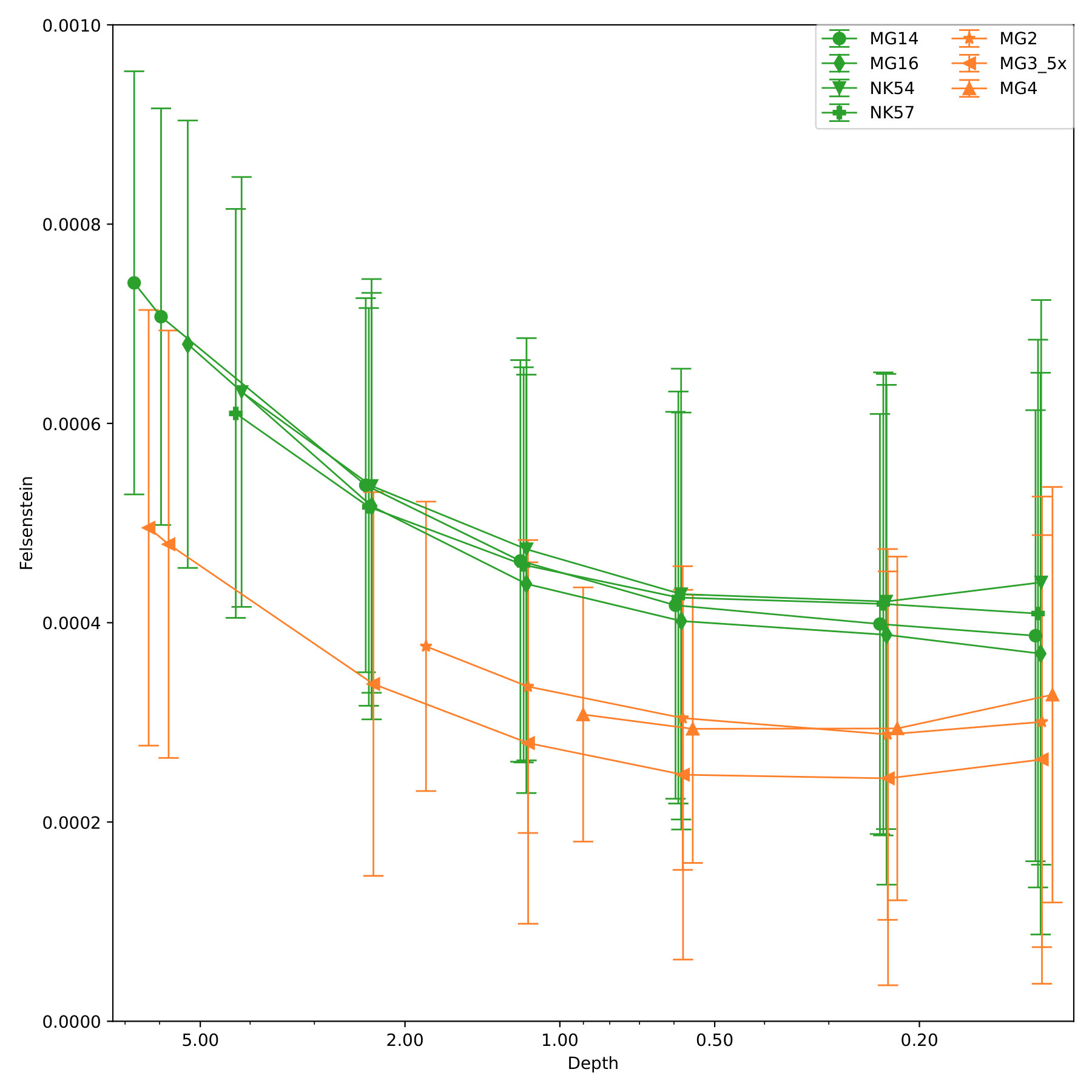

**Supplementary Figure S12: Felsenstein heterozygosity estimates calculated using a genotype likelihood framework implemented in ATLAS**. Individuals were incrementally downsampled to ensure comparability between higher and lower coverage individuals. Irish individuals are indicated in orange and German indicated in green. Error bars represent …

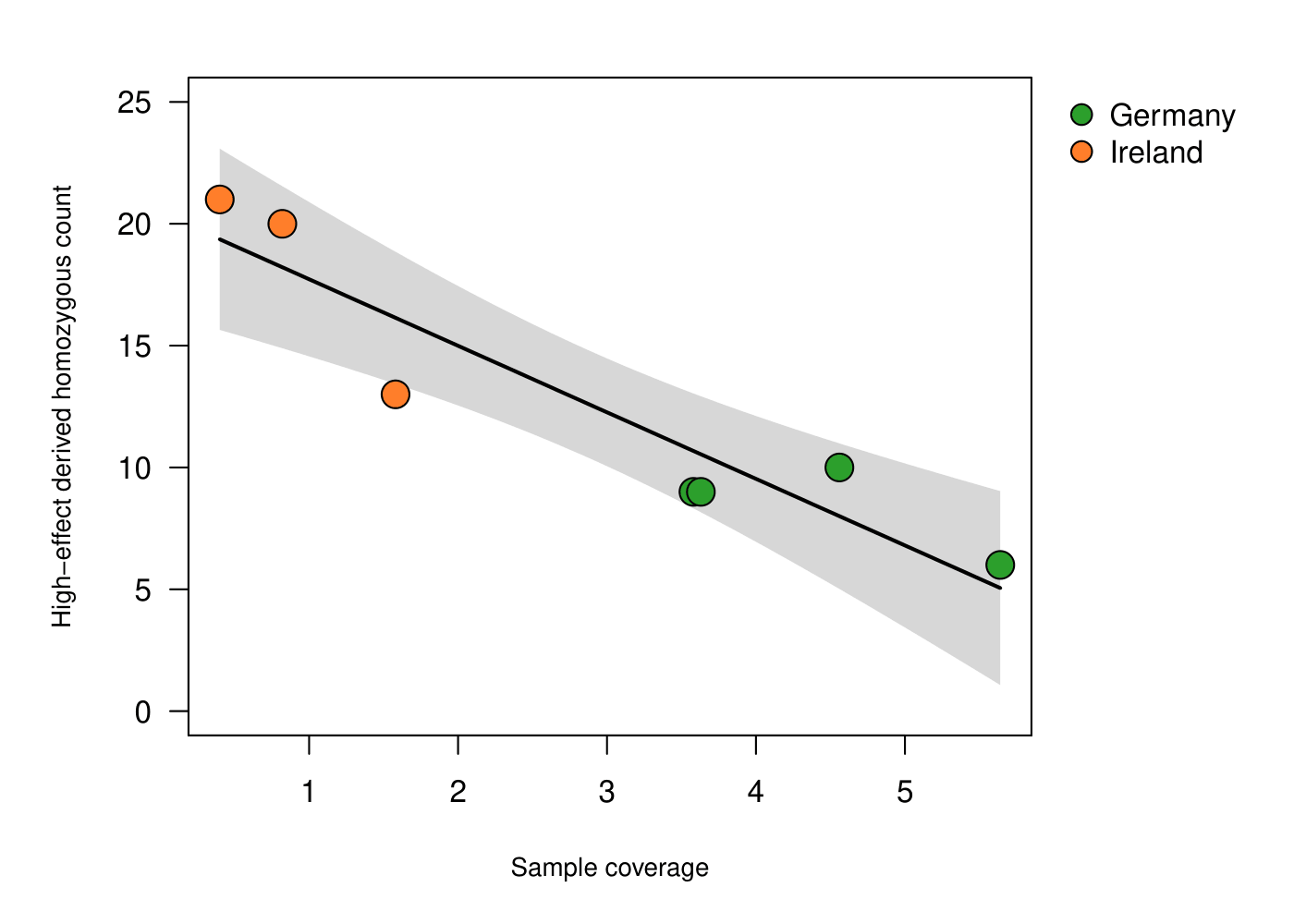

**Supplementary Figure S13. Linear regression between sample coverage and high-effect derived homozygous count across low-coverage samples from Ireland (orange) and Germany (green).** The analysis shows a significant dependence of high-effect derived homozygous count on sample coverage.

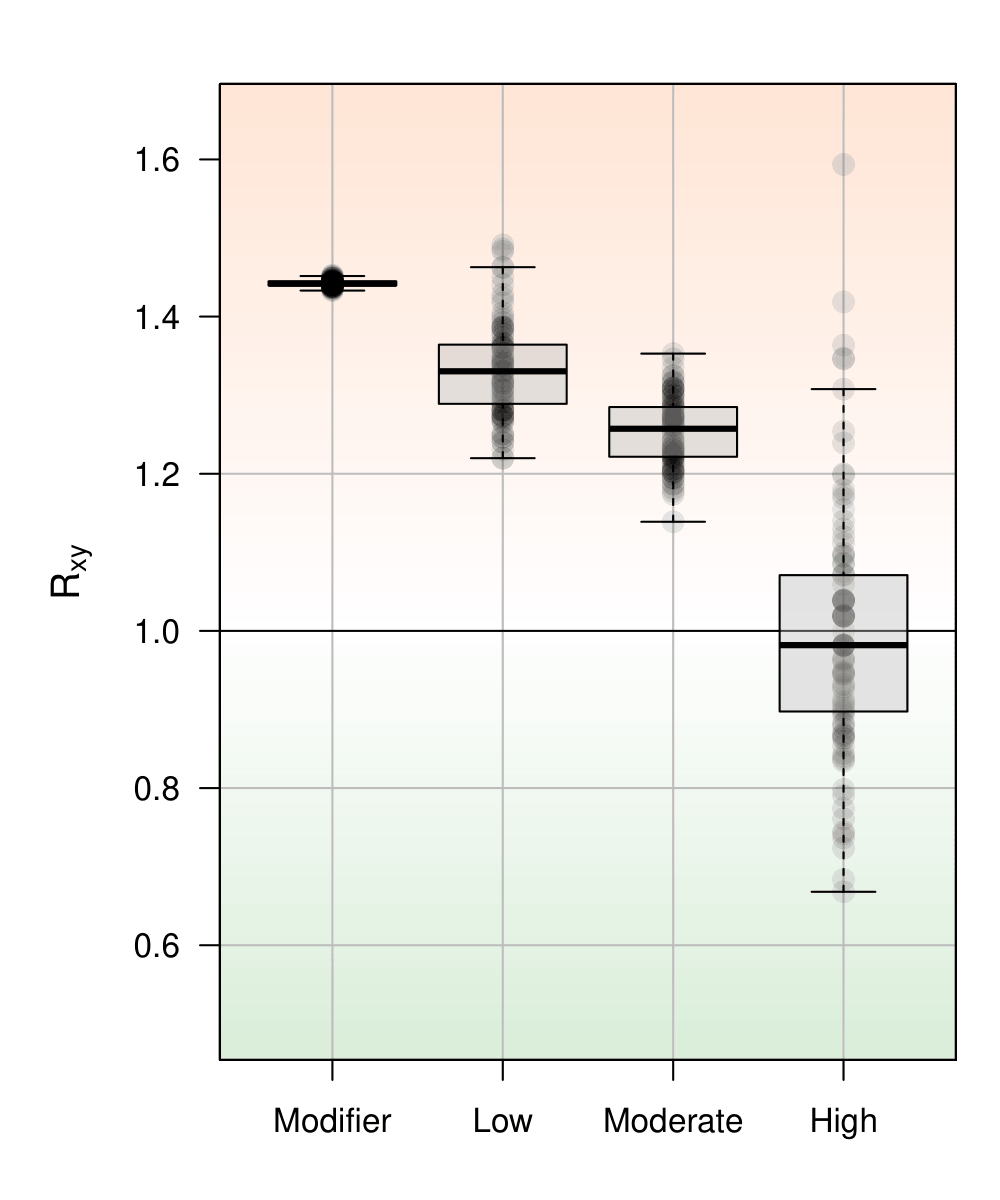

**Supplementary Figure S14. Distribution of R_xy_ values calculated between Irish (x) and German (y) populations across 100 pseudo-haploid resampled replicates.** In contrast to R’_xy_, R_xy_ is not normalized by modifier-impact sites and therefore reflects raw differences in derived allele accumulation between Irish and German populations. The color gradient highlights the direction and magnitude of the statistic, with increasing red intensity for R_xy_ > 1 (Irish enrichment) and increasing blue intensity for R_xy_ < 1 (German enrichment).

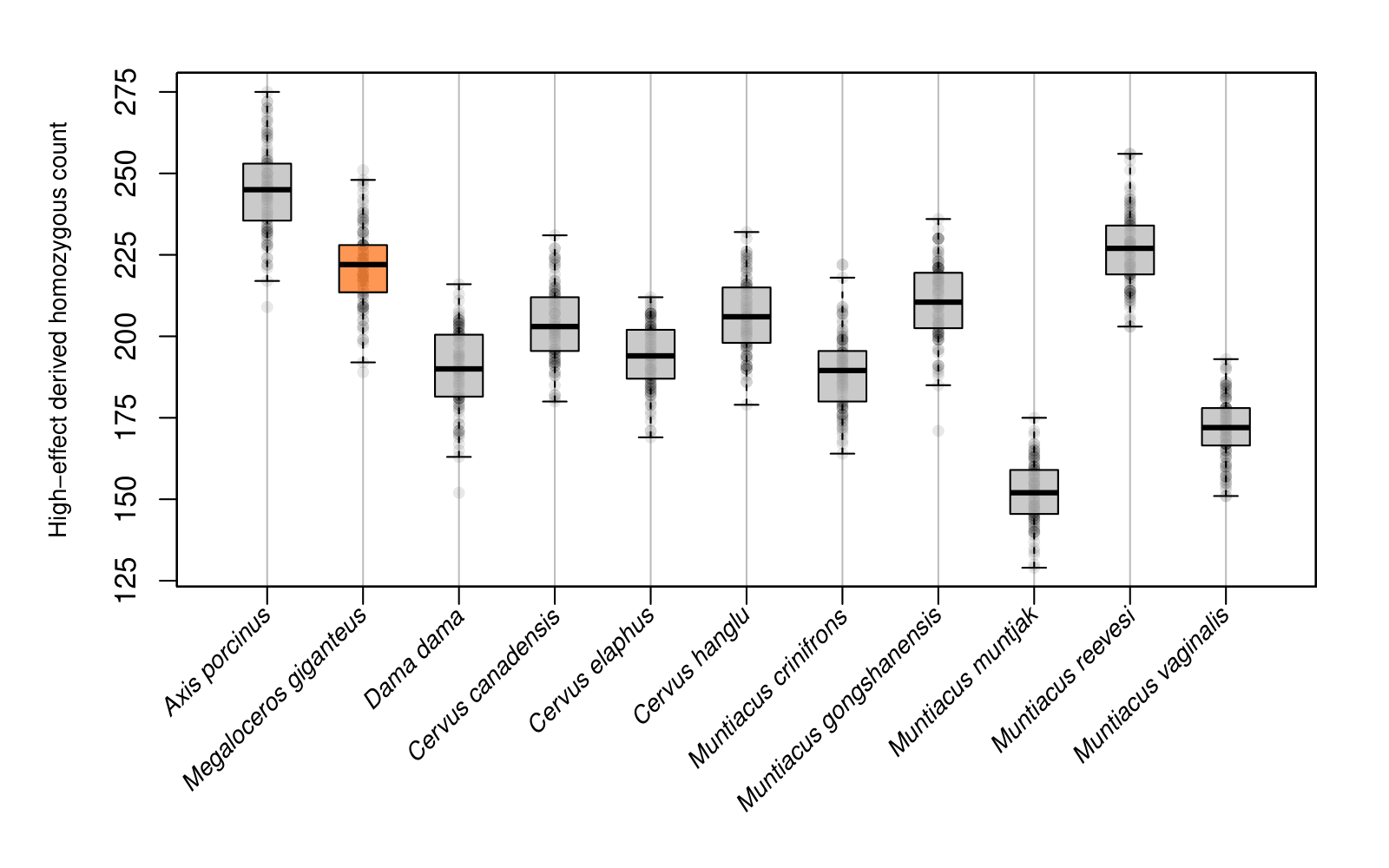

**Supplementary Figure S15. Distribution of the number of homozygous genotypes for the derived allele at high-impact sites, calculated across 100 replicate subsets of 1000 randomly sampled sites per ingroup, using only high-coverage samples.**

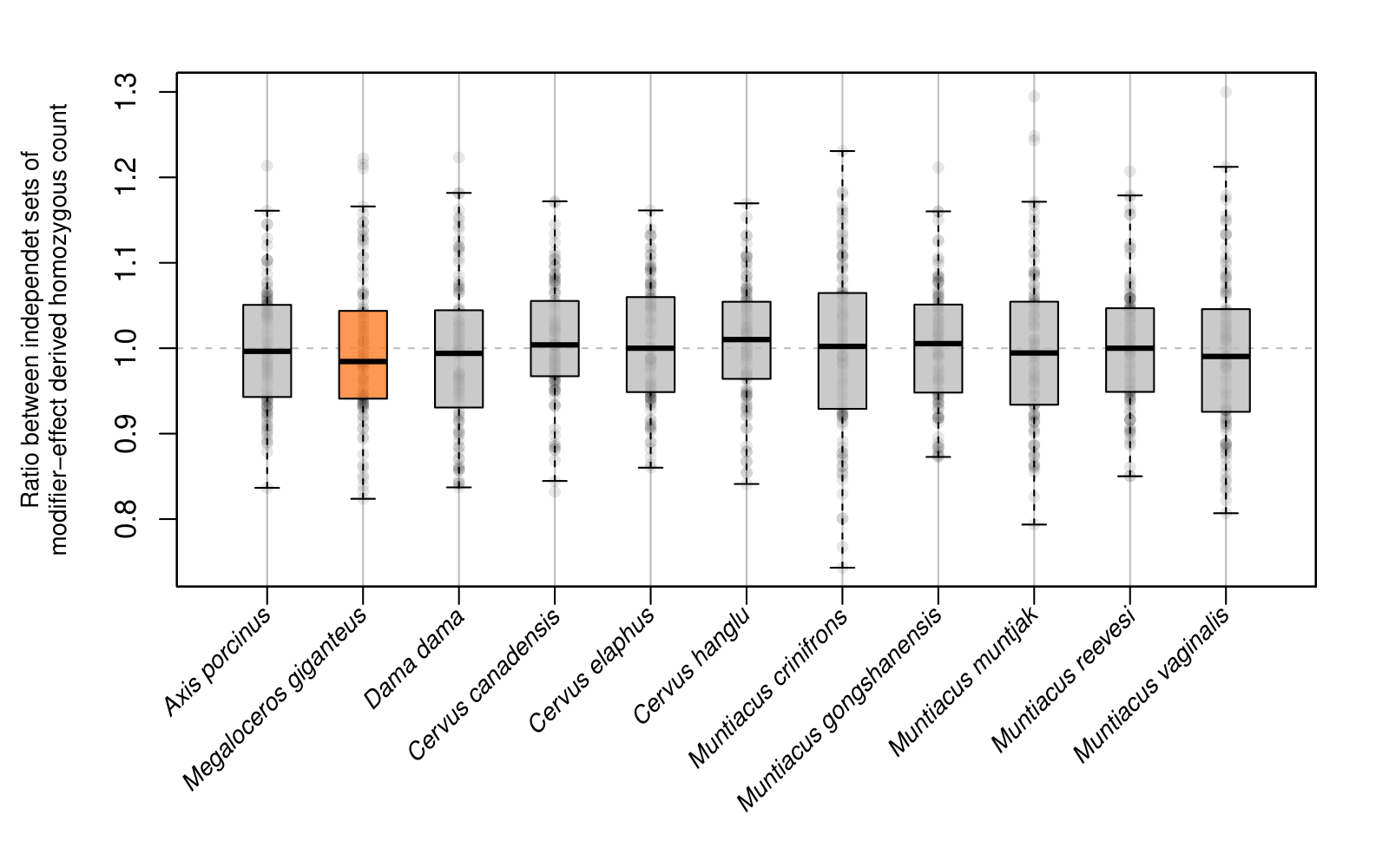

**Supplementary Figure S16.** **Validation of modifier-impact site sampling**. Distribution of the ratio between the number of homozygous genotypes for the derived allele in two independently sampled sets of modifier-impact sites, using only high-coverage samples. This analysis was performed to assess the consistency and representativeness of the sampled modifier sites. Ratios close to 1 across replicates confirm that the sampled sets are drawn from the same underlying distribution.

**Supplementary Tables**

**Supplementary Table S1: Megaloceros sample information and mapping statistics**

**Supplementary Table S2: Modern sample information and mapping statistics.** Genome ID presented if individual was mapped to a conspecific reference genome

**Supplementary Table S3: Heterozygosity and ROH estimates for all high-coverage individuals included in this study, mapped to both conspecific and heterospecific reference genomes.** “ROH adjusted for reference bias” refers to the minimum heterozygosity threshold used to define ROH, which was adjusted based on the relative difference between mappings to conspecific and heterospecific reference genomes.

**Supplementary Table S4: Raw and one-window-bridged FROH values for *Megaloceros*, *Dama dama*, and *Cervus canadensis* mapped to different reference genomes.** Under the bridging approach, single non-ROH windows flanked by ROH windows were reclassified as ROH.

**Supplementary Table S5: Heterozygosity estimates for low-coverage individuals (MG14L3, MG16L1, and MG3 downsampled to ~5×).** Estimates were calculated from sites with standardized 5× coverage (minimum and capped), and sites were considered heterozygous when at least two of five reads supported an alternative allele. Heterozygosity is reported for transversions only and for all sites. Counts of sites with one or two alternative alleles are shown for each substitution class.

**Supplementary Table S6: Genes inferred to be under episodic positive selection on the *Megaloceros* lineage based on HyPhy BUSTED analyses (P ≤ 0.05).** Gene identifiers and corresponding BUSTED P-values are shown.

**Supplementary Table S7: Significantly enriched functional terms and pathways identified using Metascape from genes under episodic positive selection.** Terms span Gene Ontology biological processes, KEGG, Reactome, and WikiPathways categories (q < 0.05). LogP values indicate enrichment significance, and associated gene symbols are listed for each term.
